# A cellular atlas of the developing meninges reveals meningeal fibroblast diversity and function

**DOI:** 10.1101/648642

**Authors:** John DeSisto, Rebecca O’Rourke, Stephanie Bonney, Hannah E. Jones, Fabien Guimiot, Kenneth L. Jones, Julie A. Siegenthaler

**Author notes:** Corresponding Author: Julie A. Siegenthaler, PhD, University of Colorado, School of Medicine, Department of Pediatrics, 12800 East 19^th^ Ave MS-8313, Aurora, CO 80045 USA.

## Abstract

The meninges, a multilayered structure that encases the CNS, is composed mostly of fibroblasts, along with vascular and immune cells. Meningeal fibroblasts are a vital source of signals that control neuronal migration and neurogenesis yet strikingly little is known about their development. We used single cell RNA sequencing to generate a cellular atlas of embryonic meningeal fibroblasts in control and *Foxc1-KO* mice in which severe CNS defects arise from failed meningeal fibroblast development. We report unique transcriptional signatures for dura, arachnoid and pial fibroblasts and identify S100a6 as the first unique marker of the pial layer. We describe a new meningeal fibroblast subtype marked by µ-Crystallin expression and show these cell types and markers are conserved in human fetal meninges. Our analysis demonstrates layer specific production of extracellular matrix components, transporter expression, and synthesis of secreted factors. Lastly, the cellular atlas of *Foxc1-KO* meninges provides insight into their severe phenotype, confirming a massive loss in arachnoid and dura fibroblasts and *Foxc1-KO* pial fibroblasts are so altered that they cluster as a different cell type based on gene expression. These studies provide an unprecedented view of meningeal fibroblast development, highlighting unexpected fibroblast diversity and function, while providing mechanistic insights into the meninges role in CNS development.

## Introduction

The meninges, made up of the dura, arachnoid and pial layers, encases the CNS from its earliest stages of development and persists as a protective covering for the adult brain. The meninges, largely composed of meningeal fibroblasts, also contains resident immune cells along with blood and lymphatic vessels. Meningeal fibroblasts have emerged as critical players in CNS development (reviewed in (Dasgupta and Jeong, 2019; Siegenthaler and Pleasure, 2011). They are the major source of basement membrane (BM) proteins that form the glial limitans (Hecht, et al., 2010a; Halfter, et al., 2002; Hartmann, et al., 1992), and meninges-derived Cxcl12 is required for migration of Cajal-Retzius cells (Borrell and Marin, 2006), proliferation of cerebellar radial glial progenitor cells (Haldipur, et al., 2015; Haldipur, et al., 2014) and proper positioning of granule cell progenitors in the hippocampus and cerebellum (Reiss, et al., 2002; Zhu, et al., 2002). Bmps produced by the meninges help control cortical layer formation (Choe and Pleasure, 2018; Choe, et al., 2012), while meningeal fibroblast production of retinoic acid directs cortical neurogenesis and cerebrovascular development (Boucherie, et al., 2018; Haushalter, et al., 2017; Mishra, et al., 2016; Siegenthaler, et al., 2009b). Much of our understanding for the role of meningeal fibroblasts in CNS development comes from studies of *Foxc1* knockout (KO) and hypomorph mice which develop severe neocortical and cerebellar defects resulting from impaired meninges formation and function (Aldinger, et al., 2009; Siegenthaler, et al., 2009b; Zarbalis, et al., 2007). *FOXC1* mutations underlie human CNS developmental defects, further underscoring its importance in CNS development (Haldipur, et al., 2017; Aldinger, et al., 2009).

Despite well-documented roles for meningeal fibroblasts in brain development, characterization of these cells is still based primarily on histological and electron microscopy studies. Further, progress on understanding meninges development has not advanced much beyond identifying the origins for meningeal fibroblasts (neural crest and mesoderm (Jiang, et al., 2002)) and some work on the timing of their developmental emergence (Siegenthaler, et al., 2009a). There is a clear need for better molecular characterization of the meningeal fibroblast populations. This will help accelerate discovery of the origins of these specialized fibroblasts, mechanisms that guide their development, and their functions in the developing and adult brain.

Here we present single cell RNA sequencing (scRNA-seq) analysis of E14.5 mouse telencephalic meningeal fibroblasts from control and *Foxc1-KO* mice. We define the transcriptional signatures of embryonic pial, arachnoid and dural fibroblasts, including identification of a first-ever pial fibroblast marker, S100a6. We present evidence for a previously unknown subtype of meningeal fibroblast we term ceiling cells, so named as the meninges are like a ‘ceiling’ for the brain, that are enriched for expression of µ-Crystallin. We show that our novel markers for the pia (S100a6), the arachnoid (CRABP2) and ceiling cells (µ-Crystallin) in mouse identify specific meningeal populations in the human fetal brain, indicating development of meningeal fibroblasts is conserved between mouse and human. ScRNA-seq and cluster analysis of the *Foxc1-KO* meninges fibroblasts revealed a near absence of pial, arachnoid and dural fibroblasts and, rather unexpectedly, a pial-like meningeal fibroblast population that clustered separately in the single cell data analysis. We believe our single cell analyses of the embryonic meningeal fibroblasts will provide a valuable resource that will aid in the study of embryonic and, possibly, the adult meningeal fibroblast populations.

## Results

### scRNA-seq and cluster analysis of embryonic meningeal fibroblasts

To carry out scRNA-sequencing analysis on developing meningeal fibroblasts, we dissected E14.5 telencephalon meninges from a *Collagen-1a1-GFP/+* (*Col1a1-GFP/+*) embryo. *Col1a1-GFP* is a well characterized transgenic mouse line and labels meningeal fibroblasts (Kelly, et al., 2016; Soderblom, et al., 2013). We chose to analyze only the telencephalic meninges as this region of the meninges is most affected in *Foxc1* mutants (Haldipur, et al., 2015; Siegenthaler, et al., 2009b). We used flow cytometry to sort GFP+ cells from the dissected telencephalic meninges of a single embryo and executed single cell capture and sequencing of 2943 cells using the 10X Genomics platform. Principal component analysis followed by t-Distributed Stochastic Neighbor Embedding (t-SNE) plotting of cells enabled cell cluster identification. Further analysis in t-SNE space and Seurat R-package analysis revealed the presence of 9 clusters (Fig. S1A). 94% (2777) of captured cells were identified as meningeal fibroblasts based on enriched expression for known meningeal fibroblasts genes (*Foxc1, Col1a1, Nr2f2, Tbx18, Zic1 and Cxcl12)* (Fig. S1B, related to Fig. 1). The remaining 6% of captured cells (166), found in four clusters, were non-fibroblasts cells known to be present in the meninges (endothelial cells, pericyte/vascular smooth muscle cells, monocytes, neural cells Fig. S1C-F, related to Fig. 1) that were inadvertently captured during the flow cytometry sort. We excluded these non-meningeal fibroblasts cells from our further analyses of the meningeal fibroblasts.

**FIGURE 1.**
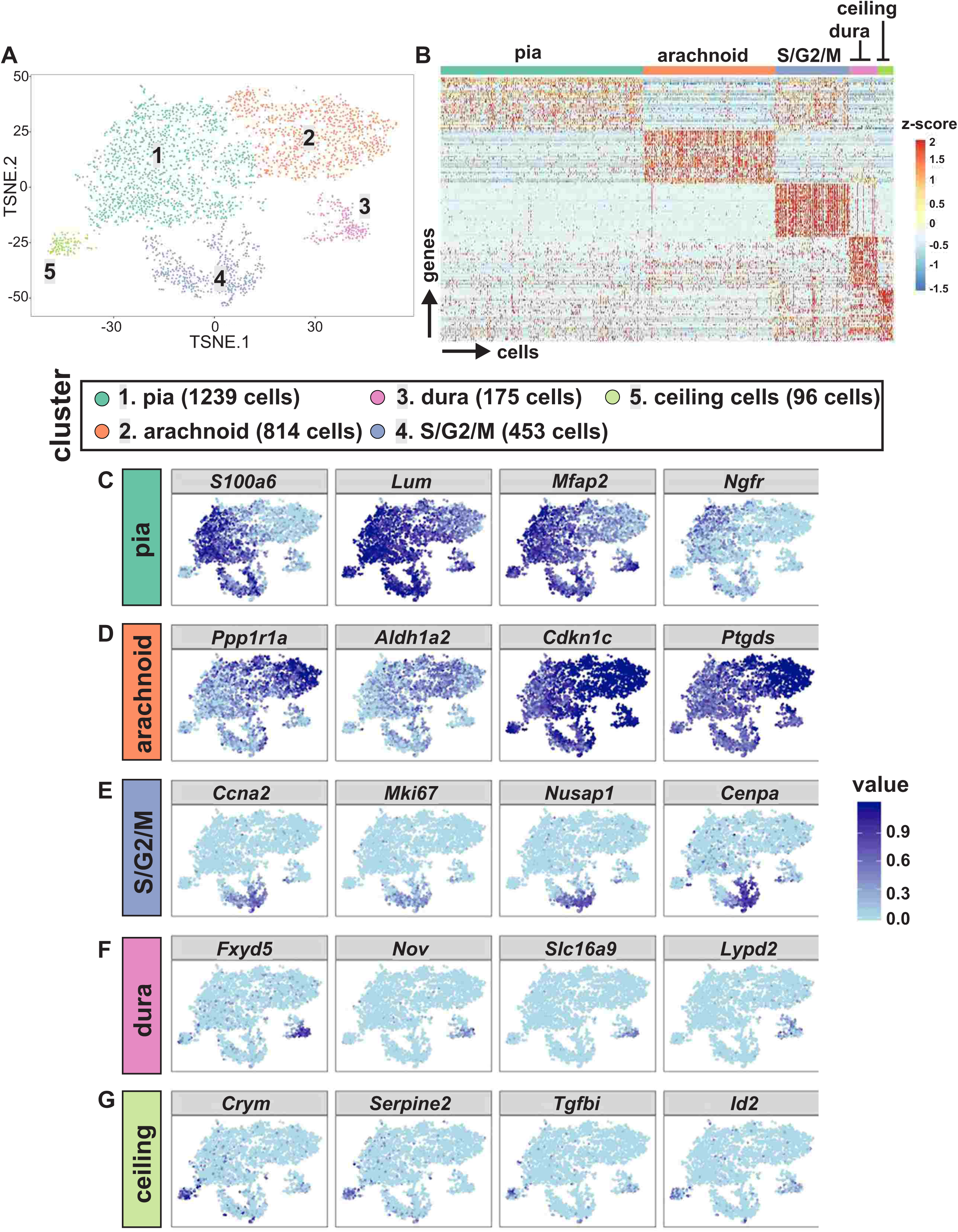
Meningeal fibroblasts cluster based on gene expression into four phenotypic groups and cycling cells. (A) Principal component analysis followed by 2D tSNE plotting of 2,777 telencephalic meningeal fibroblasts from an E14.5 *Col1a1-GFP+;Foxc1*^*+/+*^ embryo. Colors depict four meningeal fibroblast subtypes (#1 pia, #2 arachnoid, #3 dura and #5 ceiling cells) and a cluster containing cycling cells (#4). (B) Heat map showing gene expression by individual cells grouped on the basis of tSNE cluster (cells are columns and genes are rows). C – G. tSNE expression plots of individual genes with enriched expression in each of the meningeal fibroblast clusters.

The meningeal fibroblasts were found in five clusters and displayed distinct gene expression signatures within each cluster (Fig. 1A and 1B; list of genes for heat map in 1B in Supplemental data file 1). Among the five meningeal fibroblast clusters, the orange-labeled cluster (#2) showed enriched expression for the arachnoid marker *Aldh1a12* (Raldh2) (Zarbalis et al., 2007, Siegenthaler et al., 2009), while the lavender cluster (#3) was identified as dura based on enriched *Foxc2* expression (Zarbalis et al., 2007, Siegenthaler et al., 2009) (Fig 1A, B, D, F). The green cluster (#1), which contained the most cells, showed enriched expression for *S100a6, Lum, Mfpa2 and Ngfr* (Fig. 1C). We validated selective expression of S100a6 in the *Col1a1-GFP+* pial fibroblasts (detailed in Fig. 2) and therefore identified this as the pia cluster. The lime green cluster (#5), showed enriched expression of *Crym*, a gene not previously described in meningeal fibroblasts but whose protein product, µ-Crystallin, binds to thyroid hormone in the cytosol (Borel, et al., 2014). We named cells within this cluster ceiling cells. The blue cluster (#4) showed enriched expression for genes associated with actively dividing cells, specifically in S/G2/M stages of the cell cycle. This S/G2/M cluster showed expression of genes from all the meningeal subtypes (*S100a6, Mfap2, Aldh1a2, Crym, Serpine2*, Fig. 1C, D and G) indicating this cluster contains a mix of pial, arachnoid, and ceiling meningeal fibroblasts undergoing DNA synthesis or cell division. Differentially expressed genes in each control meningeal fibroblast cluster vs all other control meningeal fibroblast clusters are provided in Supplemental data file 2.

**FIGURE 2.**
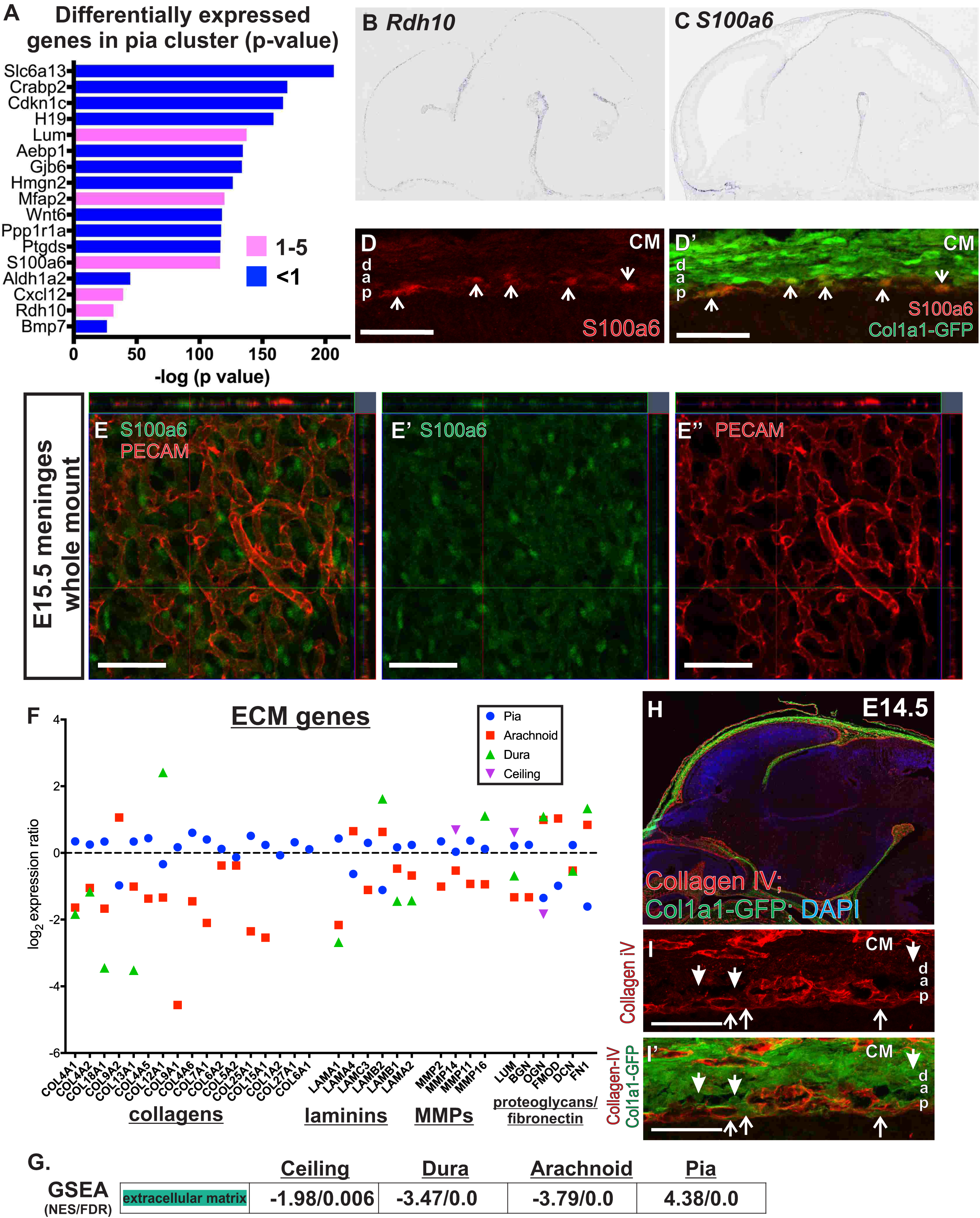
Analysis and validation of genes enriched in the pia cluster. (A) Bar graph depicts genes with greatest enrichment (ER 1-5) or depletion (ER <1) in the pial cluster. Bar length represents the negative base-10 log of p value and color represents ratio of each gene’s mean expression in the pia cluster versus all other fibroblasts (ER). (B-C) RNA in-situ hybridization showing *Rdh10* and S100a6 expression in E14.5 mouse embryo (Eurexpress.org). (D-D’) S100A6 positive cells (D, red) co-localized with *Col1a1-GFP* (D’, green) in the E14.5 telencephalic meninges (white arrows); d = dura, a = arachnoid, p = pia, CM = calvarial mesenchyme (scale bar=50μm). (E-E”) Z-stack confocal image of S100a6 (green) and PECAM (red) in E15.5 whole mount of meninges (scale bar=50μm). (F) Graph depicts log2 expression ratio for ECM genes in the pia, arachnoid, dura and ceiling cell clusters. (G) Gene set enrichment analysis (GSEA) for extracellular matrix genes in ceiling, dura, arachnoid and pia clusters reported as normalized enrichment score (NES) and false discovery rate (FDR). (H) Low magnification image of Collagen-IV expression in E14.5 *Col1a1-GFP* sagittal brain sections. (H) Higher magnification images of the telencephalic meninges depict enriched expression of Collagen-VI in GFP+ pial fibroblasts (open arrows) but not in GFP+ cells in the arachnoid and dura (closed arrows) (scale bar=50μm).

Taken together, this shows that meningeal fibroblasts from the pia, arachnoid, dura have distinct gene expression profiles, cluster as distinct subtypes, and we identify a previously unknown meningeal fibroblast subtype, ceiling cells.

### Analysis of pial fibroblasts

Pial fibroblasts are typically defined by their location at the interface between the meninges and brain tissue. No pial fibroblast specific markers have been identified and little is known about pial-specific functions in brain development. To begin to fill in these gaps, we identified genes with differential expression in the pial cluster (#1) as compared to other clusters. The gene expression ratio (ER) is the mean expression of an individual gene within a cluster divided by the mean expression of that gene in all other control fibroblast clusters. Genes with enriched expression (>1) in the pial layer include *Lum* (ER 1.15, p = 3.81×10^−138^), *Mfap2* (ER 1.27, p = 1.45×10^−120^), *Rdh10* (ER 1.38 p = 5.79×10^−40^) and *S100a6* (ER1.49, p = 4.81×10^−117^) (Fig. 2A; Supplemental data file 3). Lumican (*Lum*), microfibril-associated glycoprotein-1 (MAGP1 gene: *Mfap2*), and biglycan (*Bgn*) (Fig. S2A, related to Figure 2) are all proteoglycans and components of the extracellular matrix (ECM). *Cxcl12* (ER 1.19, p = 5.79×10^−40^), a key meninges-derived factor required for brain development, was enriched in the pial cluster (Fig. 2A) but, as shown in the *Cxcl12* t-SNE, it is also expressed in the arachnoid cluster and to a much lesser extent in the ceiling cell and dura clusters (Fig. S2A, related to Fig. 2). Some of the largest differences in expression in the pia cluster result from reduced expression (<1) of genes that are enriched in other clusters (Fig 2A), most notably *Slc6a13, Crabp2, Aldh1a2* and *Bmp7*, which are enriched in the arachnoid cluster (see Fig. 3A).

**FIGURE 3.**
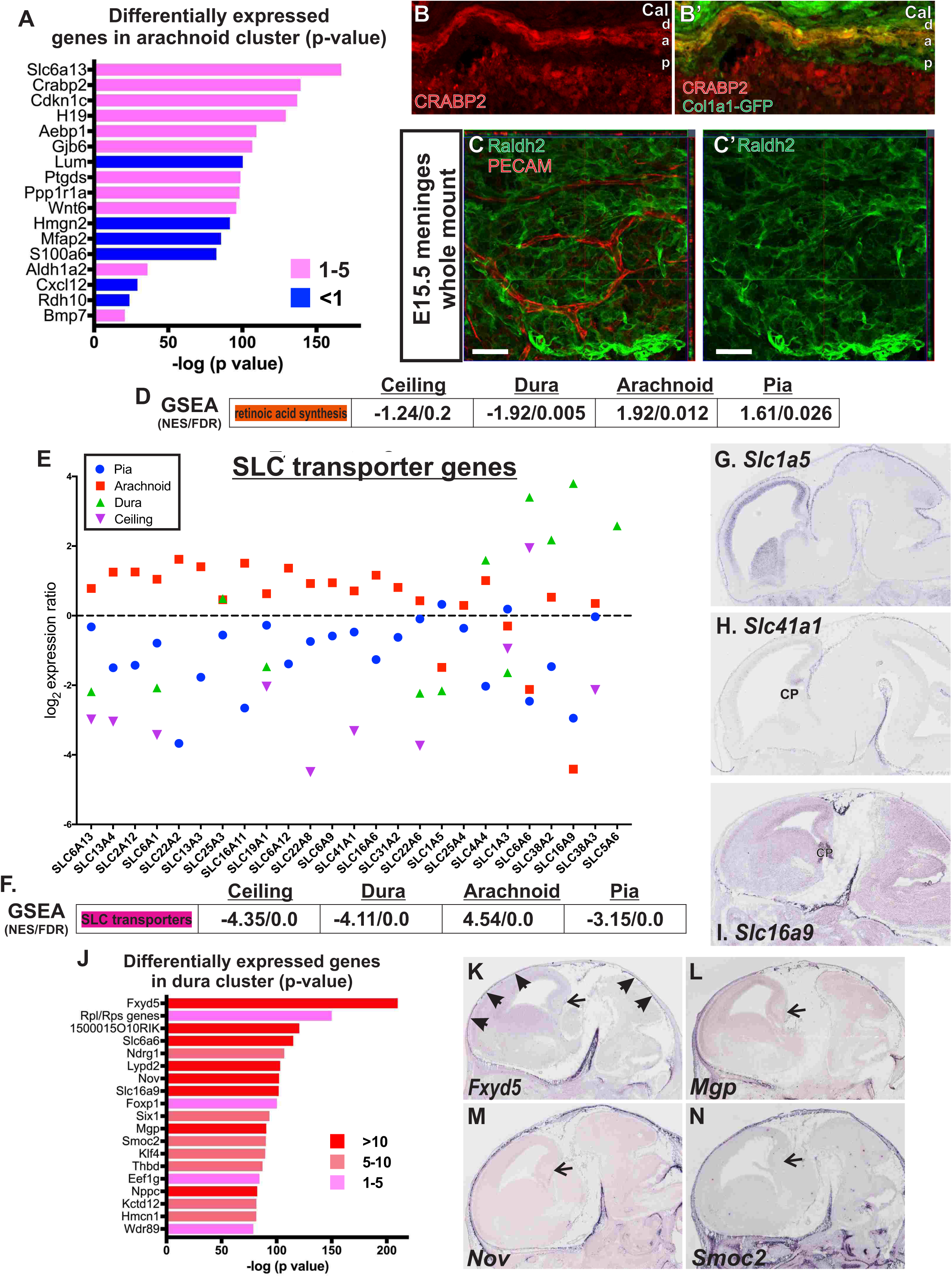
Analysis and validation of genes enriched in the arachnoid and dura cluster. (A) Bar graph depicts genes with enrichment (ER 1-5) or depletion (ER <1) in the arachnoid cluster. Bar length represents the negative base-10 log of p value and color represents ratio of each gene’s mean expression in the arachnoid cluster versus all other fibroblasts (ER). (B-B’) CRABP2 positive cells (B, red) co-localized with *Col1a1-GFP* (B’, green) in the E14.5 telencephalic meninges (white arrows); d = dura, a = arachnoid, p = pia, CM = calvarial mesenchyme (scale bar=50μm). (C-C’) Z-stack confocal image of Raldh2 (green) and PECAM (red) in E15.5 whole mount of meninges (scale bar=50μm). (D) GSEA results for genes involved in retinoic acid synthesis in ceiling, dura, arachnoid and pia clusters reporting normalized enrichment score (NES) and false discovery rate (FDR). (E) Graph depicts log2 expression ratio for SLC transporter genes in the pia, arachnoid, dura and ceiling cell clusters. (F) GSEA for SLC genes synthesis in ceiling, dura, arachnoid and pia clusters reporting NES and FDR. (G-I) RNA in-situ hybridization depicting *Slc1a5* (pia enriched, G), *Slc41a1* (arachnoid enriched, G) and *Slc16a6* (dura enriched, G) expression in E14.5 mouse embryo (Eurexpress.org). (J) Bar graph depicts genes with greatest enrichment (1-5) or depletion (<1) in the dura cluster. (K) RNA in-situ hybridization showing *Fxyd5* expression in E14.5 mouse embryo (Eurexpress.org); filled arrowheads identifies *Fxyd5* signal in the meninges overlaying the neocortex but not extending into the boundary between the telencephalon and thalamus (open arrows). (L-N) RNA in-situ hybridization showing *Nov, Mgp and Smoc* expression in E14.5 mouse embryo (Eurexpress.org).

Expression of two pia cluster-enriched genes, *Rdh10*, an enzyme required in retinoic acid biosynthesis, (Fig. 2B) and *S100a6* (Fig. 2C) was evident in the E14.5 meninges in the fore-mid- and hindbrain (*in situ* images from Eurexpress.org). Pia-selective expression of S100a6 protein expression was evident in the meninges around all brain regions at E14.5 (Fig. S2B, related to Fig. 2) and localized to *Col1a1-GFP+* cells immediately adjacent to the neocortex but was absent in GFP+ cells in the arachnoid and dura layers and the calvarial mesenchyme that covers the meninges at E14.5 (Fig. 2D). We developed a whole mount meninges immunolabeling method to better visualize the relationship between S100a6+ meningeal fibroblasts and the pia-located perineural vascular plexus (PNVP; labeled with endothelial marker PECAM). The PNVP appeared as a honeycomb pattern typical of vasculature undergoing angiogenic growth and remodeling. S100a6+ pial cell bodies were in the gaps of the honeycomb pattern in the same layer (Fig. 2E). Collectively, these data show selective localization of S100a6 to pial meningeal fibroblasts and provides strong evidence for S100a6 as a pia fibroblast marker.

The pial basement membrane (BM) is vital for normal brain development, serving as an attachment point for radial glial cells and a physical barrier to migrating neurons. Meningeal fibroblasts, in particular those of the pial layer, are the predicted source of pial BM components. To investigate pial ECM production by fibroblasts, we assembled a list of ECM-related genes (collagens, laminins, matrix metalloproteinases or MMPs, and proteoglycans) and plotted their expression in each of the four unique fibroblast clusters (Fig. 2F; Supplemental data file 3). Pial fibroblasts had the highest expression of several ECM-related genes, including *Col4a1* (ER 1.27, p = 3.43×10^−99^) and *Lama1* (ER 1.35, p = 4.33×10^−45^) two important components of the pial BM (Poschl, et al., 2004; Halfter, et al., 2002) (Fig. S2A, related to Fig. 2). This was confirmed using gene-set enrichment analysis (GSEA), demonstrating enrichment of ECM-related genes in pial meningeal fibroblasts versus the other three clusters (Fig. 2G). Of note, transcripts for certain ECM genes were enriched in other meningeal fibroblast clusters. *Col9a2* and *Col12a1* are enriched in arachnoid and dura meningeal fibroblasts, respectively, while transcripts for proteoglycans *Ogn* (osteoglycin), *Fmod* (fibromodulin), and *Fn1* (fibronectin) were enriched in both arachnoid and dura clusters but low in pia (Fig. 2F, Fig. S2C). Immunostaining analysis using a Collagen-IV (Col-IV) antibody on E14.5 brain sections from *Col1a1-GFP* mice confirm the enriched expression Col-IV by fibroblasts in the pial layer (Fig. 2H, I). Col-IV expression in in meningeal whole mounts strongly labeled the vascular BM but was also visible in adjacent pial fibroblasts (Fig. S2D, related to Fig. 2).

Collectively, our analysis of pial fibroblasts demonstrates that these cells provide key factors needed for brain development through participation in retinoic acid synthesis (based on enriched expression of *Rdh10*), a major source of Cxcl12 production, and meningeal BM proteins. Further, we provide evidence that S100a6 is a unique marker of embryonic pial fibroblasts.

### Analysis of arachnoid fibroblasts

Genes enriched in the arachnoid cluster underscore its key role in retinoic acid synthesis (*Crabp2* (ER 2.17, p = 4.73×10×10^−140^) and *Aldh1a2;* Raldh2 (ER 1.69, p = 1.17×10^−36^)) and a predicted role in transport (*Slc6a13* (ER 1.72, p = 1.90×10^−167^) (Yaguchi, et al., 2019; Zhang, et al., 2018; Yasuda, et al., 2013). Other genes with a highly enriched expression signature in arachnoid fibroblasts were the secreted glycoprotein *Angptl2* that modulates inflammation and angiogenesis (ER 2.28, 8.88×10^−97^), *Aebp1* (ER 1.99, p = 3.02×10^−112^), which encodes the extracellular matrix-associated aortic carboxypeptidase-like protein (ACLP) with a known role in fibroblast regulation (Blackburn, et al., 2018) and the Wnt ligand *Wnt6* (ER 2.17, p = 1.19 ×10) (Fig. S3A; Supplemental data file 3). *Angptl2* gene expression at E14.5 was highly enriched in the meninges (Supp. Fig. 3B, Eurexpress.org), confirming our single cell analysis.

Retinoic acid production by meningeal fibroblasts is critical to neocortical and cerebrovascular development and retinoic pathway genes *Crabp2* and *Aldh1a2* showed enriched expression in arachnoid cells. We confirmed CRABP2 expression in the E14.5 meninges using *in situ* analysis (Eurexpress.org; Fig. S3C, related to Fig. 3) and with immunostaining (Fig 3B, Fig. S3D). CRABP2 protein expression in sagittal E14.5 sections mirrored the *in situ* signal, showing expression in the meninges and in other brain regions (Fig. S3D). Immunostaining of CRABP2 in E14.5 brain sections from *Col1a1-GFP* mice showed CRABP2 expression was high in *GFP+* fibroblasts of the arachnoid and dura overlaying the neocortex but was absent from GFP+ pial meningeal fibroblasts at the pial surface (Fig. 3B). Expression in the dura layer is consistent with high expression of *Crabp2* in the dura cluster (Fig. S3A). CRABP2 was also evident in the neocortex just below the pial surface, representing radial glial endfeet that have enriched CRABP2 expression (Boucherie, et al., 2018) and, possibly, Cajal-Retzius cells that express *Crapb2* (Loo, et al., 2019). We confirmed Raldh2 (*Aldh1a2*) protein expression in arachnoid cells using whole mount immunostaining of E15.5 meninges. Raldh2+ arachnoid meningeal fibroblasts formed a sheet of cells located just above the pial-located PNVP (Fig. 3C). Rdh10 and Raldh2 are both needed to make retinoic acid however our data show that Rdh10 is expressed by the pial fibroblasts, while Raldh2 is enriched in arachnoid fibroblasts. This expression pattern is consistent with a mechanism where the pial and arachnoid fibroblasts work in conjunction to make retinoic acid, with each layer responsible for a different synthesis step. This observation is further supported by GSEA that demonstrate significant enrichment for retinoic acid synthesis genes in pial and arachnoid cluster but not in dura or ceiling clusters (Fig. 3D).

Barriers at the blood-brain (CNS endothelium) and blood-CSF interfaces (choroid plexus or CP) prevent uncontrolled intercellular movement of essential nutrients, ions and amino acids from the blood into the CNS (reviewed in (Hladky and Barrand, 2016). Because these molecules are needed for brain growth and homeostasis, transporters such as like solute carrier (SLC) influx transporter proteins are highly expressed in CNS endothelial cells and choroid plexus epithelia to ensure regulated delivery of essential molecules into the brain and CSF (reviewed in (Saunders et al., 2013). In the meninges, the arachnoid barrier layer contains tight junctions between meningeal fibroblasts cells (Nabeshima et al., 1975). Therefore, the arachnoid layer provides a selective barrier between the dura and the CSF-filled subarachnoid space and we find the arachnoid cluster transcriptome reflects this specialized function. Expression of the solute transporter *Slc6a13* (GABA transporter) was highly enriched in arachnoid cells and we identified many SLC transporter genes with enriched expression in arachnoid cells, including *Slc13a4* (sulfate transporter) and *Slc41a1* (Mg++ ion transporter) (Fig. 3E, Supplemental data file 3). GSEA using a geneset of SLC genes expressed in the meningeal fibroblast clusters confirmed enrichment of SLC genes in the arachnoid cluster (Fig. 3F). Of note, the dura cluster was uniquely enriched for expression of *Slc5a6* (biotin transporter), *Slc16a9* (monocarboxylate transporter) and *Slc38a2* (amino acid transporter) (Fig. 3E, Fig. S3E) whereas the pial cluster was enriched for *Slc1a3* (glutamate transporter) and *Slc1a5* (amino acid transporter) (Fig. 3E, Fig. S3E). In situ expression at E14.5 (Eurexpress.org) demonstrated expression of *Slc1a5* in the meninges and in the telencephalic ventricular zone that contains neural stem and progenitor cells (Fig. 3G). Arachnoid enriched gene *Slc41a1* occurred almost exclusively in the meninges (Fig. 3H); this was in contrast to dura enriched *Slc16a9*, which is highly expressed in both the meninges and the CP (Fig. 3I). Gap junctions are a feature of some barrier structures, mediating intracellular communication. Consistent with this, we noted enriched expression of gap junctional genes *Gja1* (Cx43), *Gja6* (Cx33) and *Gjb2* (Cx26) in the arachnoid cluster (Fig. S3F, related to Fig. 3) and to a lesser extent the dura cluster. Cx43 protein expression at E14.5 was evident in the meninges and the skin (Fig. S3G, related to Fig. 3). In the meninges overlying the neocortex, bright Cx43 clusters, typical of the plaques that connexins form at contacts between cells, were evident in (Fig. S3H, related to Fig. 3).

Collectively, our analyses show that arachnoid cells are enriched in transporters, confirm a critical role for arachnoid cells in retinoic acid production, and demonstrate pial and arachnoid cells likely work cooperatively to produce retinoic acid.

### Analysis of dura fibroblasts

The dura cluster differs markedly in its gene expression from arachnoid, pial and ceiling cell clusters. The dura cluster is enriched for expression of genes that encode constituents of the large and small ribosomal subunits (*Rpl* and *Rps* gene), the cell membrane ion transport regulator *Fxyd5* (ER 11.7, p = 1.3×10^−210^), and the transcription factors *Foxp1* (ER 3.73, p = 6.00×10^−101^) and *Six1* (ER 3.00, p = 3.39×10^−94^) (Fig. 3J; Supplemental data file 3). Several genes that were selectively expressed in dural fibroblasts include *Fxyd5, Foxp1, Nov* (encodes a secreted cysteine rich protein), *Smoc2* (encodes a matricellular protein), *Ndrg1* (tumor suppressor, gene mutation causes Charcot-Marie-Tooth disease type 4D) and *Kctd12* (modulates GABAB receptor kinetics, upregulated in some tumors) (Fig. S4A, related to Fig. 3). In-situ hybridization labeling *Fxyd5* expression at E14.5 (Eurexpress.org) confirms expression in a distinct layer around the telencephalon and over the hindbrain (Fig. 3K, arrowheads), but not between brain regions (arrows in Fig 3K). The dura adheres to the underside of the calvarium and therefore, unlike the pia and arachnoid layers, does not extend between brain regions. Therefore, the *Fxyd5 in situ* signal is consistent with labeling in the dura only. Dural expression patterns of the genes *Nov, Mgp* and *Smoc2* were similar to *Fxyd5* (Fig. 3L-N), validating results from scRNA-seq analysis. Collectively, these data demonstrate that, even prior to development of the calvarium, dural fibroblasts have a distinct gene expression profile and point to several new potential markers of the dura.

### Analysis of ceiling cell fibroblasts

Ceiling cells expressed a unique set of genes not expressed by other meningeal fibroblast clusters, most notably *Crym* (11.4, p = 3.01×10-93) which encodes the protein µ-Crystallin, the transcription factor *Ebf1* (5.56, p = 7.49×10^−97^), and *Serpine2* (12.4, p = 1.37×10^−145^), which encodes for the serine protease inhibitor protease nexin-1 (PN-1) (Fig. 4A, B; Supplemental data file 3). To detect ceiling cells in the embryonic meninges we examined *in situ* expression for *Crym* at E11.5 (Allan Brain Atlas) and E14.5 (Eurexpress.org). At E11.5, *Crym in situ* signal was evident in the meninges at the border between the telencephalon and the thalamus (Fig. 4C, arrow) and in the meninges in the hindbrain, above the isthmic roof plate (Fig. 4C, arrowhead). At E14.5 the Crym *in situ* signal shows a strongly labeled strip of meninges that border the future hippocampus (medial pallium/hippocampal allocortex), above the dorsal thalamus, and in the meninges around the olfactory bulb but not over the neocortex (Fig. 4D, arrows). *Crym* signal can be seen in the meninges adjacent to the basal part of the terminal hypothalamus (Fig. 4D, arrowhead), and also in neural portions of the future hippocampus and the ventricular zone of the midbrain (E11.5 and E14.5, asterisks in Fig. 4C, D). Immunostaining analysis shows a similar pattern with µ-Crystallin expressing cells in the meninges between the future hippocampus and thalamus (Fig. 4F). Higher magnification images showed µ-Crystallin+ ceiling cells adjacent to PNVP blood vessels, more typical of arachnoid than pial meningeal fibroblasts, that intermingle with the PNVP (Fig. 4G). µ-Crystallin+ cells in the meninges were *Col1a1-GFP+*, consistent with other meningeal fibroblast subtypes (Fig. 4H, I). *In situ* analysis of *Serpine2*, showed a similar pattern of expression as *Crym* with the exceptions that: (1) *Serpine2* signal was present in the meninges above the boundary between the midbrain/hindbrain (Fig. 4E, asterisk) (2) *Serpine2* signal was present in the choroid plexus (CP), and (3) no *Serpine2* signal was detected in the brain. This demonstrates that ceiling cells are a previously unknown subtype of meningeal fibroblast that does not form a distinct meningeal layer but rather localizes to specific areas of the mouse embryonic meninges.

**FIGURE 4.**
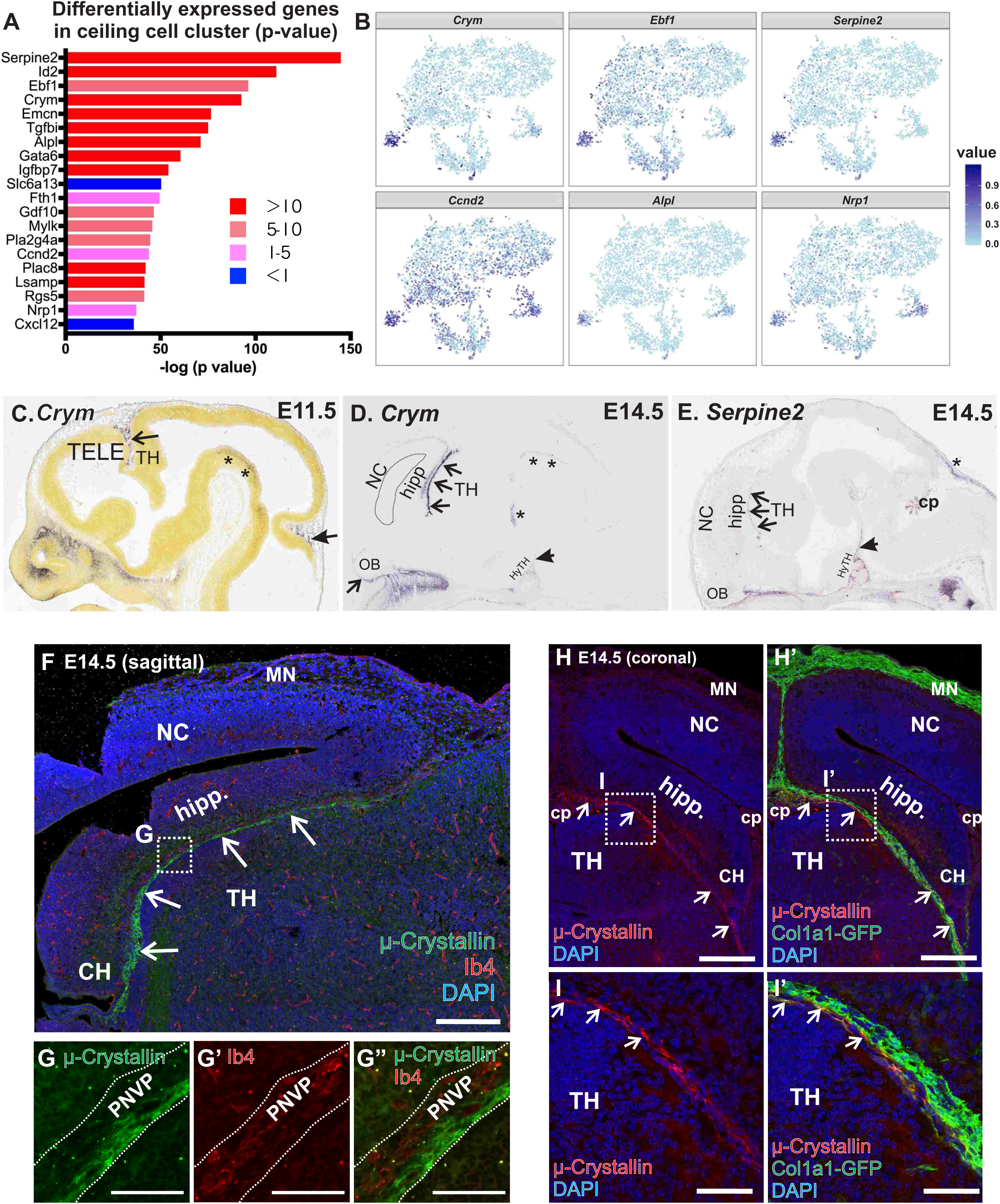
Ceiling cells are region-specific meningeal fibroblasts. (A) Bar graph depicts genes with greatest enrichment (ER 1-5) or depletion (ER <1) in the ceiling cell cluster. Bar length represents the negative base-10 log of p value and color represents ratio of each gene’s mean expression in the ceiling cell cluster versus all other fibroblasts (ER). (B) Expression plots of individual genes with enriched expression in the ceiling cell cluster shown in tSNE space. (C) RNA in-situ hybridization showing ceiling cell enriched gene *Crym* in the meninges at E12.5 (Allan Brain Atlas), located at the boundary between the telencephalon (TELE) and the thalamus (TH) (open arrow). (D) *Crym* signal in the at E14.5 (Eurexpress.org), olfactory bulbs (OB). Non-telencephalon meninges locations include near the hindbrain (E12.5, closed arrow) and adjacent to the hypothalamus (HyTH) (E14.5 closed arrow). Non-meninges area of *Crym* signal are in the hindbrain neuroepithelium (asterisks). (E) RNA in-situ hybridization showing ceiling cell enriched gene *Serpine2* in the meninges at E14.5 in similar regions as observed for Crym signal in D (open and closed arrows) and in other areas of the meninges, including overlaying the hindbrain. Serpine2 signal is also evident in the choroid plexus (CP) in the 4^th^ ventricle. (D) E14.5 saggital brain section with μ-Crystallin immunolabeling of the meninges (MN) in between the hippocampus (hipp) and thalamus (TH), this does not extend into the meninges overlaying the neocortex (NC) (scale bar=200μm). (G) Magnified area in (F) shows μ-Crystallin+ cells adjacent to IB4+ blood vessels in the perineural vascular plexus (PNVP). (H, I) E14.5 coronal brain section of *Col1a1-GFP* animal shows μ-Cystallin+ cells in between the hippocampus and thalamus are GFP+ (scale bars=200μm (H), 50μm(I)).

### Ingenuity Pathway Analysis of meningeal fibroblast clusters

To characterize gene expression within the meningeal fibroblast cell clusters, we performed Ingenuity Pathway Analysis (IPA) to identify pathways of enriched or depleted gene expression (Table 1). Dura and arachnoid clusters were strongly enriched in expression of genes in the oxidative phosphorylation pathway, indicating relatively greater metabolic levels, as compared to the pia cluster. Ceiling and dura clusters showed enrichment in vasculogenesis and angiogenesis pathway genes and correspondingly greater expression levels of genes involved in protein synthesis.

**TABLE 1:**
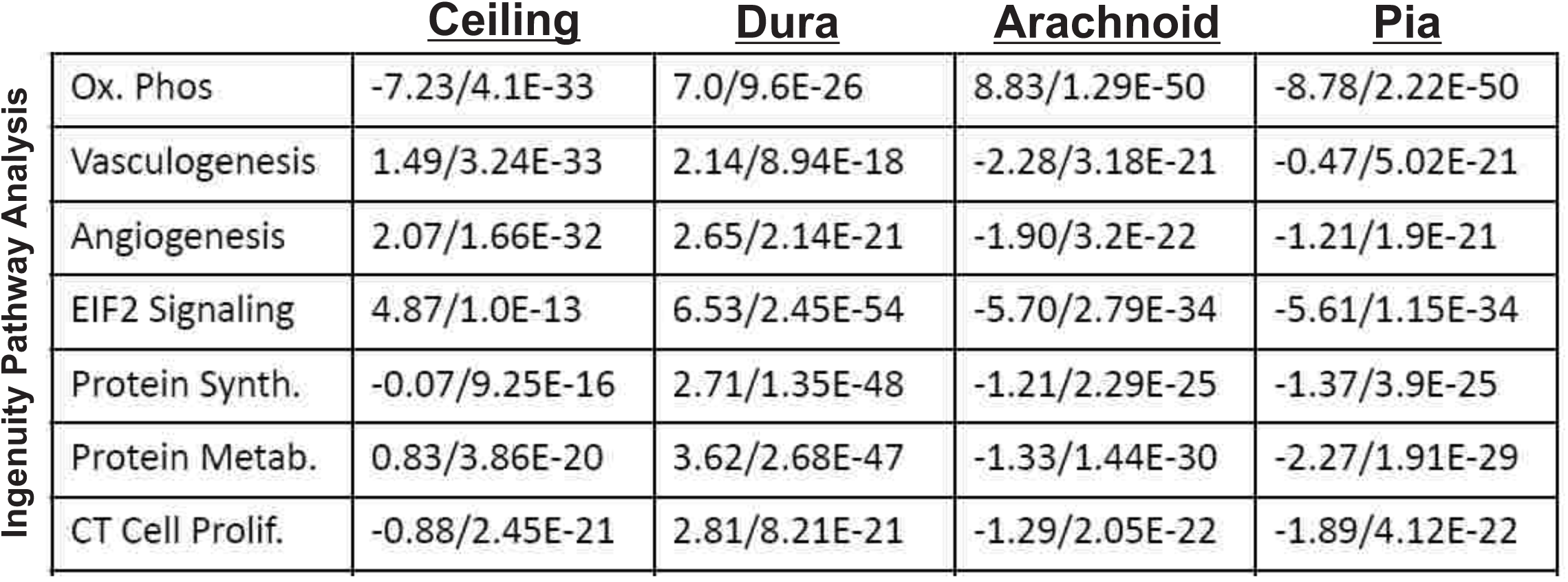
Ingenuity Pathway Analysis (IPA) of control meningeal fibroblast clusters. IPA analysis on ceiling, dura, pia and arachnoid clusters, focusing on the Biological Functions pathways that are the most relevant to developing tissues. Values presented are Z-score/p-value. Ox. Phos = oxidative phosphorylation.

| Ingenuity Pathway Analysis |  | <u>Ceiling</u> | <u>Dura</u> | <u>Arachnoid</u> | <u>Pia</u> |
| --- | --- | --- | --- | --- | --- |
|  | Ox. Phos | -7.23/4.1E-33 | 7.0/9.6E-26 | 8.83/1.29E-50 | -8.78/2.22E-50 |
|  | Vasculogenesis | 1.49/3.24E-33 | 2.14/8.94E-18 | -2.28/3.18E-21 | -0.47/5.02E-21 |
|  | Angiogenesis | 2.07/1.66E-32 | 2.65/2.14E-21 | -1.90/3.2E-22 | -1.21/1.9E-21 |
|  | EIF2 Signaling | 4.87/1.0E-13 | 6.53/2.45E-54 | -5.70/2.79E-34 | -5.61/1.15E-34 |
|  | Protein Synth. | -0.07/9.25E-16 | 2.71/1.35E-48 | -1.21/2.29E-25 | -1.37/3.9E-25 |
|  | Protein Metab. | 0.83/3.86E-20 | 3.62/2.68E-47 | -1.33/1.44E-30 | -2.27/1.91E-29 |
|  | CT Cell Prolif. | -0.88/2.45E-21 | 2.81/8.21E-21 | -1.29/2.05E-22 | -1.89/4.12E-22 |

### Conservation of mouse meningeal fibroblast subtype markers in the human fetal meninges

To determine whether meningeal fibroblast subtype specific markers are conserved between mouse and human developing meninges, we examined expression of CRABP2 (arachnoid/dura in mouse), S100a6 (pia in mouse), and µ-Crystallin (ceiling cells in mouse) in the meninges overlying the neocortex in human fetal brain (19 gestational week or GW). We examined two areas of the fetal cortical meninges, one overlaying the frontal cortex (region A) and the other in the sylvian sulcus overlaying the insular cortex (region B) (Fig. 5A). In region A, CRABP2 was expressed by two morphologically distinct meningeal cell layers, a loose network of CRABP2+ cells (arrows in 5B) and a compact, outer layer that seemed to contain only CRABP2+ cells (arrowheads in 5B). In region B, the meninges were much thicker and the CRABP2+ layer of loosely packed cells was considerably expanded (Fig. 5E). Immediately below this was a cell sparse region containing long strips of tissue, some of which contained CRABP2+ cells (Fig. 5E, arrows). These are the arachnoid trabeculae that traverse the subarachnoid space (SAS) in higher vertebrate and primate meninges (Mortazavi, et al., 2018). Arachnoid trabeculae and the SAS were not apparent in the meninges overlying the frontal cortex at this stage of development. CRABP2+ cells were not detected in the meningeal cells near the brain in either region. Collectively this provides evidence that CRABP2 is expressed by arachnoid cells in the developing human brain.

**FIGURE 5.**
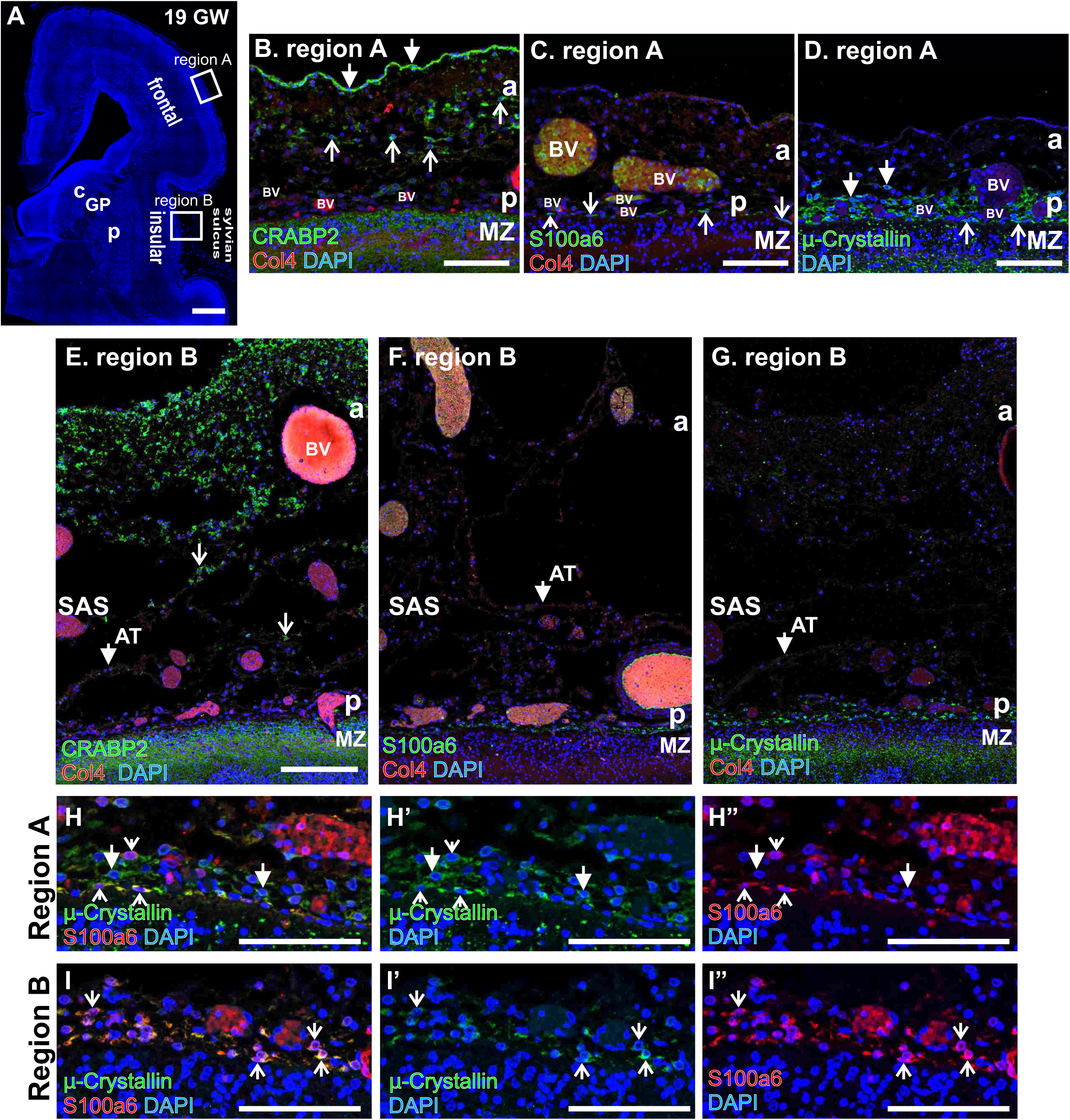
Meningeal fibroblast subtypes are present in human fetal meninges. (A) Coronal section of a 19 gestational week (GW) human telencephalon, region A is meninges overlaying the frontal cortex and region B is in the sylvian sulcus overlying insular cortex (C=caudate, GP=globus pallidus, and P=putamen). (B-D) In region A, arachnoid marker CRABP2 labels loosely (open arrows) and tightly packed cells (closed) in the meninges distal from the brain, consistent with CRABP2 labeling arachnoid fibroblasts (a=arachnoid). Pial marker S100a6+ cells were limited to a single layer immediately adjacent to the brain, consistent with being in the pia (p) (open arrows) next to Col4+ blood vessels (BV), many filled with auto-fluorescing red blood cells. μ-Crystallin+ cells were also in the pial layer, immediately adjacent to the brain surface (open arrow) and slightly distal (closed arrows). (E-G) Region B contained a much-expanded meninges with a distinct subarachnoid space (SAS), recognized by arachnoid trabeculae (AT) (arrowheads in E-G). The arachnoid layer and some trabeculae contained CRABP2+ cells (E). S100a6+ (F) and μ-Crystallin+ (G) cells were limited to the pia adjacent to the brain surface in region B. (H, I) S100a6+/μ-Crystallin+ pial cells (open arrows) and μ-Crystallin+/S100a6-(closed arrows) in regions A (H) and B (I).

S100a6+ cells were limited to cells immediately adjacent to the brain in regions A and B, intermingled with Col4+ blood vessels (Fig. 5C, arrows, and 5F), consistent with S100a6 being expressed by pial meningeal fibroblasts in the fetal human brain. µ-Crystallin+ cells, which marks ceiling cells in mouse, were observed throughout the meninges covering the fetal human cortex. This is in contrast to mouse where µ-Crystallin+ ceiling cells are not uniformly distributed (Fig. 4). The µ-Crystallin+ cells were located at the pial surface (Fig. 5D, 5G). Many µ-Crystallin+ cells at the pial surface expressed S100a6 (Fig. 5H & I, arrows), however µ-Crystallin+ cells were detected in the cell layer just above the S100a6+ layer, and these cells were S100a6 negative (Fig. 5H, arrowheads). We examined expression of CRABP2 and µ-Crystallin but did not observe co-localization in region A or B (Fig. S4B, C, related to Fig. 5), indicating CRABP2 and µ-Crystallin label distinct meningeal subtypes in developing human tissue. This data shows that arachnoid and pial markers are conserved between mouse and humans and shows that µ-Crystallin+ meningeal fibroblasts are present in human developing meninges.

### Single cell transcriptome analysis of the Foxc1-KO meningeal fibroblasts

Simultaneous with our single cell analysis of telencephalic GFP+ meningeal fibroblasts from an E14.5 *Col1a1-GFP/+; Foxc1*^*+/+*^ embryo (Fig. 1-4), we captured GFP+ cells from an E14.5 *Col1a1-GFP/+; Foxc1-KO* littermate, in which Foxc1 expression is lost globally, resulting in severely impaired meningeal and neocortical development (Siegenthaler, et al., 2009b; Kume, et al., 1998). Simultaneous cluster analysis of control and *Foxc1-KO* cells identified 12 distinct clusters (Fig. S5A, related to Fig. 6). Four clusters contained pericyte/vSMCs (mural cells), endothelial, monocytes, and neural cells; these made up only a small percentage and were captured inadvertently (Fig. S5A, related to Fig. 6). A small number of *Foxc1-KO* cells colocalized in tSNE space with the five control meningeal fibroblasts clusters described in detail in Figs. 1-4 (Fig. S5B, C, related to Fig. 6). Three clusters (#10, #11, #12) contained only *Foxc1-KO* cells (Fig. S5A-C, related to Fig. 6). Of the clusters composed only of *Foxc1-KO* cells, cluster #10 showed expression of meningeal fibroblast markers (detailed in Fig. 6) (Fig S5A-C, related to Fig. 6). Clusters #11 and #12 contained the majority of *Foxc1-KO* cells and had a gene expression signature consistent with osteogenic cell types, including *Osr2, Twist1, Twist2* and members of the Iraqoius homeobox (IRX) protein gene family (*Irx1, Irx5*) that are not detected in any of the control and *Foxc1-KO* meningeal fibroblast clusters (Fig. S5D, related to Fig. 6). We believe the appearance of these osteogenic clusters in the *Foxc1-KO* capture is a result of difficulties in dissection. Specifically, the calvarial mesenchyme, which expresses *Col1a1-GFP*, cannot be cleanly dissected away from the hypoplastic meninges of *Foxc1-KO* mice. Of note, there were also 20 *Foxc1-KO* cells that could not be classified (Fig. S5C, related to Fig. 6). Because we sought to specifically investigate the phenotype of *Foxc1-KO* meningeal fibroblasts, the two osteogenic clusters and clusters containing pericytes/vSMCs, endothelial cells, monocytes and neural cells were not considered further in our comparison of control and *Foxc1-KO* cells.

**FIGURE 6.**
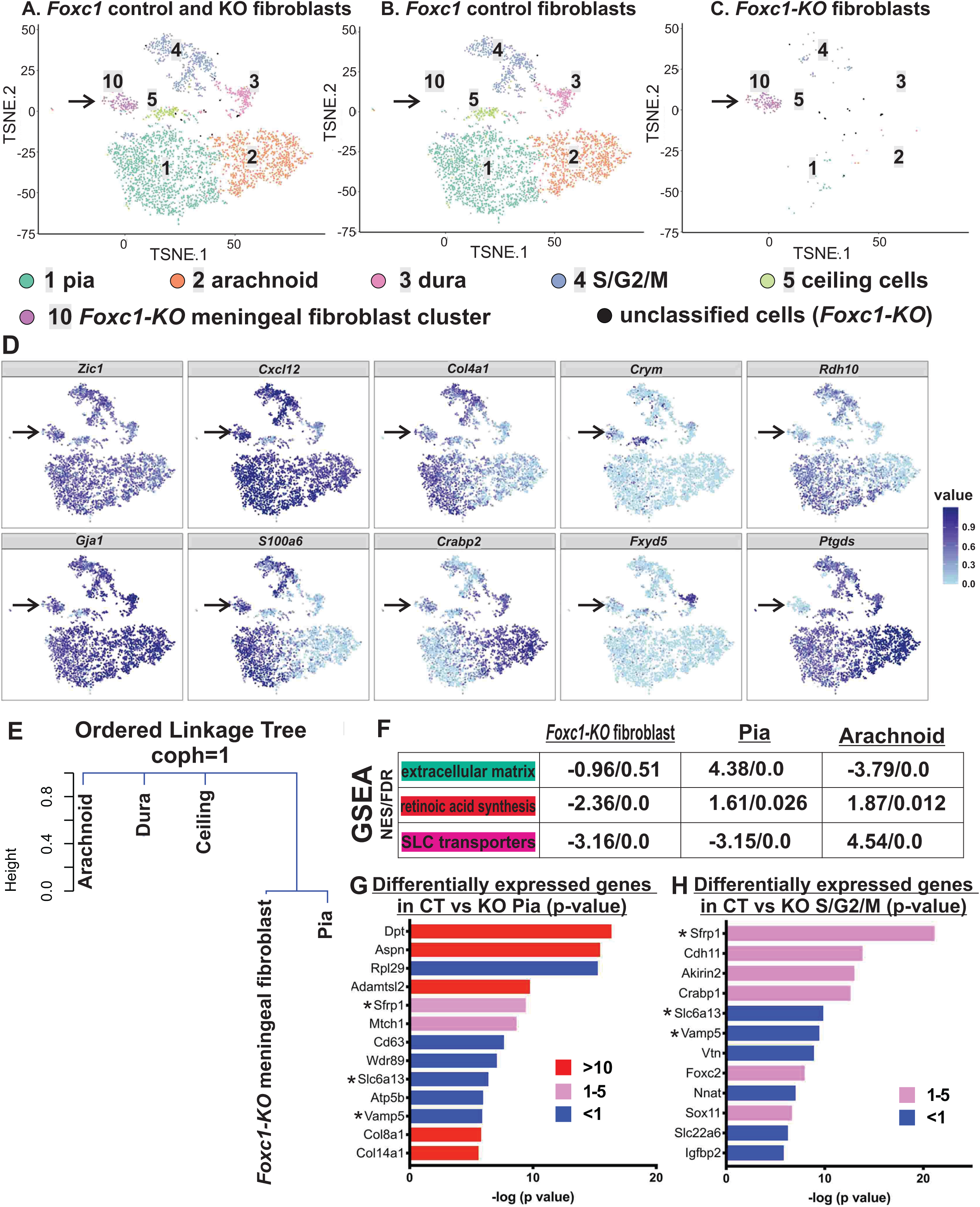
Comparative analyses of control and *Foxc1-KO* meningeal fibroblast clusters. (A-C) tSNE plots of meningeal fibroblast clusters (cluster #1-5, 10) containing control and *Foxc1-KO* cells. (D) tSNE gene expression plots depicting control and *Foxc1-KO* meningeal fibroblast clusters. Arrow indicates cluster #10, a meningeal fibroblast cluster that contains only *Foxc1-KO* cells. (E) Hierarchical clustering analysis of control meningeal fibroblast clusters and Foxc1-KO meningeal fibroblast cluster #10 demonstrates this cluster is most similar to the control pia. (F) GSEA of meninges-relevant pathways (ECM production, retinoic acid synthesis and SLC transport) in *Foxc1-KO* meningeal fibroblast cluster (#10); control pia and arachnoid values are provided for comparison (NES = normalized enrichment score and FDR=false discovery rate). (G, H) Bar graph depicts genes with enrichment (1-10, >10) or depletion (<1) in *Foxc1-KO* vs control pia and S/G2/M. Bar length represents the negative base-10 log of p value and color represents ratio of each gene’s mean expression in the *Foxc1-KO* pia or S/G2/M vs versus control pia or S/G2/M (ER). * indicate genes that are similarly enriched or depleted in *Foxc1-KO* pia and S/G2/M cells.

t-SNE plots displaying meningeal fibroblast clusters from control and *Foxc1-KO* fibroblasts, show no *Foxc1-KO* dura or ceiling cells and illustrate that most of *Foxc1-KO* meningeal fibroblasts occupy a single cluster (#10) distinct from control clusters (Fig. 6A-C, arrow). The *Foxc1-KO* cluster (#10) shows enriched expression for some common meningeal fibroblast markers (*Cxcl12, Zic1, Gja1)* (Fig. 6D), and shows enriched expression of pial markers *S100a6* and *Col4a1,* but not arachnoid/dural marker *Crabp2*, dural marker *Fxyd5* or ceiling cell maker *Crym* (Fig. 6D). Of note, *Rdh10*, which is enriched in the control pia cluster (ER 1.38, p = 2.48×10^−32^), was significantly lower in the *Foxc1-KO* cluster and *Ptgds*, expressed in all control clusters and enriched in arachnoid clusters, was nearly absent (ER 0.016, p = 3.31×10^−10^) (Fig. 6D). To understand the relationship between the *Foxc1-KO* meningeal fibroblast cluster (#10) and control meningeal fibroblast clusters, we performed a non-negative matrix factorization (NMF) hierarchical clustering analysis (Brunet, et al., 2004) and found that the *Foxc1-KO* meningeal fibroblast cluster most closely resembles the control pial cluster (Fig. 6E). The ordered consensus matrix (Fig. S6C) and cophenetic correlation value (coph=1) for this analysis show that the optimal number groups was k=4and that the clustering analysis attained a high degree of confidence. Gene expression analysis of this meningeal fibroblast *Foxc1-KO* cluster (#10) revealed enriched expression of WNT antagonist *Sfrp1*, transcription factor *Jund*, actin binding protein Profillin-2 (*Pfn2*), retinoic acid binding protein *Crabp1* and Notch inhibitor *Dlk1* (Fig. S6A, B, arrows; Supplemental data file 3). Using GSEA, we found that the *Foxc1-KO* meningeal fibroblast cluster was deficient in expression of both ECM and retinoic acid synthesis pathway genes (Fig. 6F). These analyses suggest that most *Foxc1-KO* meningeal fibroblasts are a pial-like cell population that lack critical functionality such as ECM production and retinoic acid synthesis.

A small percentage of *Foxc1-KO* cells clustered with control meningeal fibroblast clusters. The number of *Foxc1-KO* arachnoid cells (8 cells) was insufficient for meaningful comparative analysis therefore we focused on the *Foxc1-KO* pial and S/G2/M clusters. We compared gene expression between control and *Foxc1-KO* pial clusters and identified >50 genes that were significantly different between the two genotypes; a curated list of these is shown in Fig. 6G (full list in Supplemental data files 4). Similar analysis of S2/G2/M cluster comparing control and *Foxc1-KO* yielded many of the same genes enriched in the *Foxc1-KO* pial-like cluster (#10) (*Sfrp1,Crabp1*) and downregulated in *Foxc1-KO* pial cluster (*Vamp5, Slc6a13*), indicating that proliferating *Foxc1-KO* pia and pial-like meningeal fibroblasts are in this cluster (Fig. 6H, full list in Supplemental data files 3, 4). Collectively, these data show that the few meningeal fibroblasts in *Foxc1-KO* mice that still cluster with control meningeal fibroblasts are nonetheless distinctly altered in their gene expression.

### Development of all telencephalic meningeal fibroblast layers is impaired in Foxc1-KO mice

We and others have previously shown that telencephalic meninges layer development is impaired in *Foxc1-KO* mice. However, we wanted to apply our new markers and whole mount staining to better characterize the meningeal layer defects. Low-magnification images of CRABP2, (arachnoid/dura), Cx43 (in all meningeal fibroblasts enriched in arachnoid/dura), and S100a6 (pia) in E14.5 control (*Col1a1-GFP; Foxc1*^*+/+*^) and *Col1a1-GFP; Foxc1-KO* sections at the level of telencephalon demonstrate that strong expression of meningeal fibroblasts subtype markers ends ventrally in the mutants (Fig. 7B, H, N, arrowheads). High magnification images of CRAPB2 immunostaining in an area over the neocortex, illustrate the absence of strongly CRABP2+ arachnoid/dural cells in the *Foxc1-KO* mutant (Fig. 7C, D). In contrast, a CRABP2+ meningeal layer, though thinner, was detected at the base of the *Foxc1-KO* telencephalon (Fig. 7E, F). A similar pattern was observed for Cx43, with near absence of strong Cx43+ cells overlaying the neocortex in the *Foxc1-KO* mutant (Fig. 7I, J) and an attenuated Cx43+ meningeal layer at the base (Fig 7K, L). S100a6+ cells were observed in the *Foxc1-KO* mutant overlaying the dorsal neocortex however the population appeared less numerous (Fig. 7O, P, arrowheads). At the base of the telencephalon a relatively normal appearing S100a6 pial layer was apparent in *Foxc1-KO* sample (Fig. 7Q, R). Images presented in Fig. 7C-F, I-L and O-R also contain *Col1a1-GFP* signal to identify fibroblasts; these are provided in Fig. S7.

**FIGURE 7.**
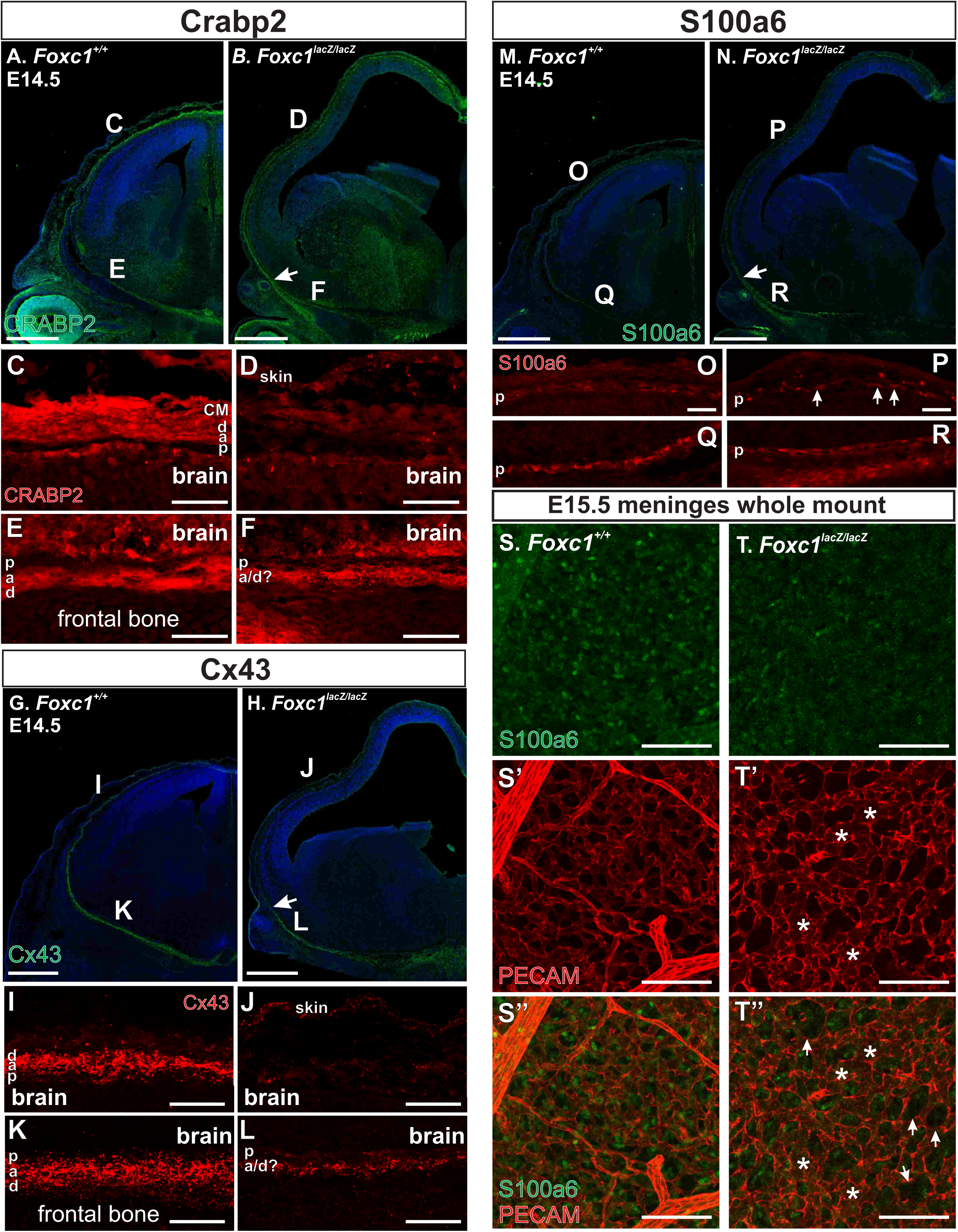
Meningeal layer development defects in *Foxc1-KO* animals. *Foxc1*^*+/+*^ and *Foxc1-KO* at the level of the telencephalon depicting CRABP2 (A, B), Cx43 (G, H) and S100a6 (M, N) expression. Arrows indicate termination of distinct meningeal marker labeling. (C-F) Higher magnification images of areas indicated in A, B of CRABP2 labeling in the meninges (arachnoid and dural) above the neocortex (C, D) and adjacent to the ventral telencephalon (E, F) in control and *Foxc1-KO* embryos. CRABP2 signal is also observed in the brain (I-L). Higher magnification images of areas indicated in I, J of Cx43 labeling in the meninges above the neocortex (I, J) and adjacent to the ventral telencephalon (K, L) in control and *Foxc1-KO* embryos. (O-R) Higher magnification images of areas indicated in Q, R of S100a6 labeling in the pia above the neocortex (O, P) and adjacent to the ventral telencephalon (Q, R) in control and *Foxc1-KO* embryos. Arrows in “P” indicated S100a6+ cells adjacent to the brain in *Foxc1-KO* sample. (S-T) E15.5 meningeal whole mount depict numerous S100a6+ pial fibroblasts interspersed with PECAM+ PNVP in control sample (S) but far fewer S100a6+ cells in the *Foxc1-KO* sample, including several areas devoid of S100a6+ cells (arrows in T”). The PNVP was dysplastic in *Foxc1-KO* samples, indicated by broad, poorly refined vasculature (asterisks in T’, T”). Scale bars = 500 μm (A, B, G, H, M, N), 100 μm (C-F, I-L, O-R) and 200 μm (S, T). p=pia, a=arachnoid, d=dura, cm=calvarial mesenchyme.

The sparseness of the S100a6+ layer in the *Foxc1-KO* mutant as compared to control was evident in E15.5 meninges whole mounts (Fig. 7S, T). In the control sample, S100a6+ cells were readily observed between PECAM+ vessels of the PNVP (Fig. 7S). In contrast, S100a6+ cells were sparse in the *Foxc1-KO* and gaps in the vasculature sometimes contained no S100a6+ cells (Fig. 7T”, arrowheads). The PNVP vasculature was notably different in *Foxc1-KO* mutant, with broad, immature vessels (Fig. 7T’, T” asterisks). Collectively, these data confirm meningeal fibroblast development is severely impaired in *Foxc1* mutant mice and the few, drastically altered meningeal fibroblasts that are present cluster as a unique cell type with pia-like characteristics.

## Discussion

Here we present the first comprehensive analyses of embryonic meningeal fibroblasts, a cell type with important functions in brain development. We have: 1) identified new meningeal fibroblast subtype markers and show these are conserved in developing human meninges; 2) generated layer specific insights into embryonic meninges functions (retinoic acid synthesis, ECM production, transport); 3) identified a previously unknown meningeal fibroblast subtype, ceiling cells; 4) and shown that all meningeal fibroblast subtypes are severely reduced in number in *Foxc1-KO* mice, replaced by a fibroblast population with pial-like characteristics.

Meningeal fibroblasts are a known source of ECM proteins that comprise the BM (Beggs, et al., 2003; Sievers, et al., 1994); our data provide important information as to which subtype(s) is producing these BM-located proteins. We show a number of ECM genes, including collagens, laminins, MMPs and proteoglycans have enriched expression in pial meningeal fibroblasts as compared with arachnoid, dura and ceiling cells. Some of these are known components of the pial BM which serve as an essential boundary segregating the brain from the meninges (Halfter, et al., 2002). We also show that key ECM genes, including laminin genes *Lama4, Lamb1* and *Lamb2*, proteoglycans *Ogn* and *Fmod*, and fibronectin (*Fn1*) have greater enrichment in dura or arachnoid cells as compared to pia. Fibronectin and laminins are key components of the pial BM; our data shows that all subtypes likely contribute to pial BM make-up. Also, it is highly likely that ECM produced by arachnoid, dural and ceiling cells have functions beyond pial BM formation. One potential example of this is enrichment of *Ogn*, the gene that encodes the secreted proteoglycan osteoglycin that has been implicated in regulation of bone formation (Lee, et al., 2018). *Ogn* has enriched expression in arachnoid and dura that are adjacent to the calvarial mesenchyme that is actively undergoing inter-membranous ossification at E14.5. The meninges, in particular the dura, have previously been shown to play a role in regulating calvarial development (Dasgupta and Jeong, 2019; Greenwald, et al., 2000; Rice, et al., 2000; Mehrara, et al., 1999); however, the meningeal fibroblast subtypes and the signals involved are incompletely understood. Our profiling of the meningeal fibroblasts during a critical period of calvarial development could be quite valuable for understanding meninges-calvarial signaling.

The meninges and the choroid plexus contain cell layers that serve as a barrier between the periphery and the CSF. Efflux and influx transporters that control the flow of solutes into and out of the CSF have been relatively well characterized in the choroid plexus (Saunders, et al., 2013; Ek, et al., 2010). Much less is known about transporter expression in the meninges, and in particular, which transporters are expressed by arachnoid barrier cells and therefore participate in solute transport into and out of the CSF (Yaguchi, et al., 2019; Zhang, et al., 2018; Yasuda, et al., 2013). We used our single cell data to shed light on the subtype specific expression of genes in the SLC transporter superfamily which consists of membrane-bound passive and active transporters from 55 different gene families that perform substrate-specific membrane transport but share minimal evolutionary commonality (He, et al., 2009). Molecules transported by SLC superfamily members include amino acids, sugars, inorganic ions, vitamins, fatty acids and essential metals. We show that cells in the arachnoid cluster are enriched in SLC genes as a group, but a few SLC genes are enriched in non-arachnoid cells. The SLC genes enriched in non-arachnoid cells have duplicate substrates to those enriched in the arachnoid layer suggesting this difference may have minimal functional significance (e.g., SLC6A6, enriched in the dura, and SLC6A13, enriched in the arachnoid, both include taurine as a substrate; and SLC16A9, enriched in the dura, and SLC16A6 and 11, enriched in the arachnoid, are all carriers for monocarboxylates such as pyruvate and lactate). Arachnoid barrier cells are a subset of arachnoid cells that form tight junctions in the adult (Nabeshima, et al., 1975), but it’s not clear when these cells appear or arachnoid barrier characteristics emerge during development. Therefore we cannot specifically link expression of SLC transporters to arachnoid barrier cells, which may emerge later than the time point we are studying, using our profiling data. However, the enrichment of SLC genes as a group in arachnoid cells supports the idea that the arachnoid layer plays an important role in controlling CSF composition through solute transport. More studies are needed on developing and mature meningeal populations to understand arachnoid barrier emergence and function and its role in regulating movement of solutes into and out of the CSF; this has important implications physiologically as well as for the delivery of therapeutic agents into the brain (Weller, et al., 2018).

Retinoic acid synthesis is an important meninges function during brain development, in particular for the neocortex. Retinoic acid biosynthesis occurs in two steps, the reversible and rate-limiting synthesis of retinaldehyde from retinol catalyzed by retinol dehydrogenase, Rdh10, and the irreversible conversion of retinal to retinoic acid catalyzed by one of the retinal dehydrogenases, encoded by *Aldh1a1, Aldh1a2* or *Aldh1a3* (Napoli, 2012). Although previous work has identified the embryonic leptomeningeal expression of *Rdh10* and *Aldh1a2* as a source of retinoic acid for the developing brain (Siegenthaler, et al., 2009b; Romand, et al., 2008; Cammas, et al., 2007; Li, et al., 2000), the precise roles of meningeal subtypes in retinoic acid biosynthesis, utilization and excretion have not been defined. While our data do not conclusively provide this information, new observations about meningeal subtype-specific roles with regard to retinoic acid can be derived from our results. In particular, pial cells have enriched expression of *Rdh10* and therefore appear to be a main source for retinaldehyde production by the meninges whereas *Aldh1a2*-enriched arachnoid cells are responsible for retinoic acid production. Thus it appears there is a previously unknown ‘division of labor’ for retinoic acid synthesis by different cells of the leptomeninges. Possibly, this is required to control retinoic acid delivery to the brain as the retinoic acid-producing arachnoid cells are separated from the brain by pia meningeal fibroblasts and vasculature. Arachnoid and dura also highly express CRABP2, an intracellular transporter that binds with high affinity to cytoplasmic retinoic acid and delivers it to retinoic acid receptors (RARs) (Napoli, 2017). GSEA did not show a significant increase in arachnoid expression of genes whose transcription is mediated by RARs (data not shown), suggesting CRAPB2 may not facilitate RAR-activated gene expression. However, CRABP2 may be promoting important non-genomic functions of retinoic acid and RARs in arachnoid and dura meningeal fibroblasts. More studies are needed to understand the function of local, meningeal retinoic acid signaling and meningeal retinoic acid synthesis; these will be greatly aided by our embryonic single cell profiling data.

Beyond providing important new insight into meningeal fibroblast function, our single cell profiling revealed, rather unexpectedly, a regionally restricted meningeal fibroblast subtype. Ceiling cells represent a unique population, distinguishable from other meningeal fibroblasts by high expression of *Crym* (Crystallin-µ) and *Serpine2* (PN-1) and their presence in distinct locations within the embryonic meninges. Our analyses so far indicate ceiling cells are bone fide meningeal fibroblasts since they express pan-meningeal fibroblast genes (*Col1a1, Zic1, Foxc1, Nr2f2*, and *Cx43*). Based on GSEA of meninges relevant functions, ceiling cells do not share retinoic acid and ECM production functions with pial and arachnoid subtypes and, in this way, they are similar to dura meningeal fibroblasts. IPA indicates potential roles in vasculogenesis and angiogenesis, a feature shared with dura. The most intriguing question, of course is, what is their function? Possibly, they have region-specific functions related to CNS development or act locally to regulate development of meninges-located cell types and structures (immune cells or blood vasculature) contributing to specialization of these areas of the meninges. Ceiling cells uniquely express PN-1 (*Serpine2*), a serine protease inhibitor that can inhibit angiogenesis (Selbonne, et al., 2012) and *Serpine2*^*−/−*^ mice display hypervascularity in the developing retina and muscle (Selbonne, et al., 2015). They also are enriched for *Nrp1* which encodes for Neuropilin-1, the receptor for the repulsive guidance cue SEMA3 ligands and VEGF-A ligand receptor on endothelial cells, acting as a co-receptor for VEGFR2. High gene expression of angiogenesis modulators indicate ceiling cells may act to locally regulate development of the pial vasculature though more studies are needed to test this and other potential functions of ceiling cells in CNS development.

Our studies of human fetal meninges show subtype expression of certain markers are conserved between mouse and human (pia: S100a6 and arachnoid: CRABP2). This is an important finding as it indicates meningeal fibroblast specification and development is likely similar between mouse and human and important functions of the meningeal fibroblasts in development, like secretion of signals that act on progenitors and migrating neurons, are also likely conserved. Consistent with this idea, studies have identified similar CNS developmental defects in *Foxc1* mouse mutants and human patients with *FOXC1* point mutations and deletions (Haldipur, et al., 2017; Aldinger, et al., 2009). The widespread distribution of µ-Crystallin + cells in the pial layer of human fetal meninges overlying the neocortex was quite different than mouse, where *Crym*/Crystallin-µ is limited to ceiling cells that localize to specific regions of the embryonic meninges. At this point, we cannot say if µ-Crystallin-+ cells in the human fetal meninges are analogous to ceiling cells in the mouse. However, our data do suggest greater cellular diversity in human fetal pial meningeal fibroblasts. In particular, we observe µ-Crystallin+ meningeal fibroblasts expressing S100a6 and negative for S100a6 overlying the cortex, implying there are potentially two subtypes in the developing human pia. While more work is needed to test this idea, it is tempting to speculate that meningeal fibroblast diversity evolved in parallel with the CNS to support the unique needs of forming a much larger and more complex human brain.

Our analysis of meningeal fibroblasts from *Foxc1-KO* mice using scRNAseq provides a more complete picture of the meninges defect in these mutants. Perhaps most important is the clear impairment in the development of pia meningeal fibroblasts in *Foxc1-KO*. We and others have assumed that the telencephalic pia is preserved in *Foxc1-KO* and the primary defects are in the formation of the arachnoid and dural layers. This assumption was based on 1) the presence of some pan-meningeal marker+ cells (Zic1, Cx43, Pdgfrβ) intermingled with the PNVP overlaying the dorsal telencephalon; and 2) the relatively well preserved pial BM in *Foxc1-KO* mice at E14.5, significant alterations in BM integrity are not seen until later in prenatal development (Hecht, et al., 2010b). Further, prior work lacked the benefit of pial specific markers to differentiate pial meningeal fibroblasts from other subtypes. We show that all meningeal fibroblasts subtypes are severely diminished in the *Foxc1-KO* mice and a pial-like meningeal fibroblast cluster emerges that is enriched for a unique set of genes. We believe that these *Foxc1-KO* pial-like cells are the S100a6+/Col1a1-GFP+ cells that sparsely occupy the pial layer overlaying the mutant telencephalon and that ‘normal’ pial meningeal fibrolasts stop at the same location as arachnoid and dura populations, at the base of the telencephalon. The *Foxc1-KO* pial-like cells appear to be poor substitutes for pial meningeal fibroblasts. This is suggested by reduced gene expressed for ECM components and components of the retinoic acid synthesis pathway, in particular *Rdh10*. Many questions remain about the *Foxc1-KO* pial-like population, including their lineage relationship to other meningeal fibroblasts, why they lack functional features, and how the absence of *Foxc1* permits their emergence. Rather disappointingly, the very low number of other meningeal fibroblast subtypes limited or prevented meaningful analysis. *Foxc1* hypomorph mutants that have milder meningeal defects (Zarbalis et al., 2007, Siegenthaler et al., 2009) is an approach we can take in future single cell studies to understand how lack of Foxc1 impairs the formation of the telencephalic meninges layers.

In summary, our analysis provides much needed characterization of fibroblasts that make up the embryonic meninges. This data set will serve as an important resource for researchers studying meninges-related structure, process, malformation and disease during development and in the adult. Further, it will serve as an important tool to identify the mechanisms that direct fibroblast diversity and specialization in the meninges and help accelerate discovery of new meningeal fibroblast subtype-specific functions.

## Materials and methods

### Animals

All mice were housed in specific-pathogen-free facilities approved by AALAC and were handled in accordance with protocols approved by the IACUC committee on animal research at the University of Colorado, Anschutz Medical Campus. The following mouse lines were used in this study: *Foxc1*^*lacZ*^ (Kume, et al., 1998) and *Col1a1-GFP* (Yata, et al., 2003). *Foxc1*^*lacZ/+*^;*Col1a1-GFP*^*GFP/+*^ male and female mice were interbred to generate *Foxc1*^*lacZ/lazZ*^;*Col1a1-GFP*^*+/GFP*^. *Col1a1-GFP*^*GFP/+*^;*Foxc1*^*+/+*^ and *Col1a1-GFP*^*GFP/+*^;*Foxc1*^*lacZ/+*^ littermates were used as controls in the scRNA-seq experiment and tissue staining.

### Sample preparation

Meningeal fibroblasts were derived from a single E14.5 *Col1a1-GFP*^*GFP/+*^;*Foxc1*^*+/+*^ and single littermate *Foxc1*^*lacZ/lazZ*^;*Col1a1-GFP*^*+/GFP*^ embryo. Using a dissection microscope, the skin and calvarial mesenchyme were removed exposing the meninges, which were then dissected from the telencephalon. GFP+ cells were isolated from the meningeal tissue by first incubating the tissue in 1.5 mL of digestion solution consisting of HBSS with Ca^2+^/Mg^2+^ (Invitrogen), 2% w/v bovine serum albumin (Calbiochem), 1% w/v glucose, 5 mg/mL Type II collagenase (Worthington) and 2% DNase, for 15 minutes at 37°C. The tissue was then triturated until well dispersed in the digestion solution. Liberation of the cells from the tissue into a single cell suspension was confirmed by viewing samples of the solution under a microscope, with additional incubation time at 37°C if needed to completely disaggregate the cells. Fluorescently-activated cell sorting (MoFlo XDP cell sorter; Beckman Coulter) was then performed using the *Col1a1-GFP*^*+/GFP*^ marker to isolate GFP+ fibroblasts from other cell types, with gating optimized to maximize GFP+ cell recovery.

### Single cell capture, library preparation and sequencing

To capture, label, and generate transcriptome libraries of individual cells, we used the 10X genomics Chromium Single Cell 3’ Library and Gel Bead Kit v2 (Cat #PN-120237) following the manufacture’s protocols. Briefly, the single cell suspension, RT PCR master mix, gel beads and partitioning oil were loaded into a Single Cell A Chip 10 genomics chip, placed into the Chromium controller, and the Chromium single cell A program was run to generate GEMs (Gel Bead-In-EMulsion) that contain RT-PCR enzymes, cell lysates and primers for sequencing, barcoding, and poly-DT sequences. GEMs are then transferred to PCR tubes and the RT-PCR reaction is run to generate barcoded single cell identified cDNA. Barcoded cDNA is used to make sequencing libraries for sequencing analysis. Sequencing was performed on an Illumina NovaSeq 6000 using paired end 150 cycle 2×150 reads. Cell capture, library prep and sequencing was performed by the Genomics and Microarray core at the University of Colorado, Anschutz Medical Campus.

### Analysis of single cell RNA-Seq data

Cellranger (2.0.2) [1] count module was used for alignment, filtering, barcode counting and UMI counting of the single cell FASTQs, the aggr module used for normalizing samples to same sequencing depth, and the reanalyze module used for final determination of gene expression and t-SNE 2D and 3D coordinates with parameters = [max_clusters=20, t-SNE_max_dim=3, num_principal_comp=20, t-SNE_perplexity=50, t-SNE_theta=0.25, t-SNE_max_iter=10000, t-SNE_stop_lying_iter=1000 and t-SNE_mom_switch_iter=1000]. A cluster of 429 cells with consistently low read counts (<30,000 per cell) was removed and the remaining cells re-analyzed with the same parameters as above. A 3D t-SNE plot was generated using plotly (2.0.15) (Plotly Technologies Inc. Collaborative data science. Montréal, QC, 2015. https://plot.ly) in a custom script and cells clustered manually based on the 3D t-SNE plot. 2D t-SNE cluster plots were generated with CellrangerRkit (2.0.0) visualize_clusters module with clustering based on the manual clustering using the 3D t-SNE plots, and gene expression plots were generated with the visualize_gene_markers module [limits=c(0,1.2)] from the log of the normalized gene expressions for each cell. A one-way analysis of variance using a linear model of gene expression ∼ sample was calculated to determine the p-value for each expressed gene in all cells in given control or *Foxc1-KO* cluster compared to its expression in all other cells (Supplemental data file 1) or to compare gene expression in *Foxc1-KO* cells verses control cells in the pia and S/G2/M clusters (Supplemental data files 2 and 4).

In order to further characterize gene expression among the meningeal fibroblast clusters identified through the t-SNE clustering analysis, we computed gene expression ratio (ER) and p-values by comparing mean expression by gene in each meningeal fibroblast cluster to the mean expression of the remaining meningeal fibroblast clusters. Supplemental data file 3 reports these data for control and *Foxc1-KO* clusters. We conducted pathway analyses using Ingenuity Pathway Analysis (IPA, Qiagen), focusing on the Biological Functions pathways as the most relevant. We performed GSEA (Broad Institute (Subramanian, et al., 2005)) using custom-prepared genesets consisting of: (i) ECM genes (collagens, laminins, matrix metallo-proteinases, proteoglycans and fibronectin); (ii) solute carriers genes (SLC genes); and (iii) genes involved in retinoic acid synthesis. In order to characterize the overall pattern of gene expression in derived from the *Foxc1-KO* cluster #10, we performed hierarchical clustering using Pearson distance, K-means clustering and non-negative matrix factorization (NMF) on the combined control and mutant meningeal fibroblast populations. Input data were mean expression by t-SNE-identified subgroup. Only NMF was effective in resolving the various subgroups.

To assign *Foxc1-KO* cells to meningeal fibroblast subtype cluster, we used t-SNE space and Seurat (v2.3.4) (Butler, et al., 2018). For most cells, the meningeal fibroblast subtype cluster determined by Seurat was consistent with its mapping in t-SNE space. If the cell was in the S/G2/M t-SNE space but not assigned to this cluster by Seurat, if the expression levels of *Ccna2, Mki67, Nusap1, Cenpa, Birc5, H2afz, Cks1b, Cks2, Lmnb1 and Stmn1* were similar to *Foxc1*^*+/+*^ S/G2/M cells, it was assigned to the S/G2/M cluster. Otherwise, it was labelled “undetermined”. None of the *Foxc1-KO* cells in the dura t-SNE space had expression levels of *Fxyd5, Nov, Slcl6a9, Lypd2, Foxp1, Smoc2, Ndrg1*, and *Kctd12* similar to *Foxc1*^*+/+*^ dura cells and were all labeled “undetermined”. None of the *Foxc1-KO* cells in the ceiling t-SNE space had expression levels of *Crym, Serpine2, Tgfbi, Id2, Ebf1, Ccnd2, Alpl, and Nrpl* similar to *Foxc1*^*+/+*^ ceiling cells and were all labeled “undetermined”. A few of the *Foxc1-KO* cells in the *Foxc1-KO* pia-like cluster were assigned by Seurat to the pia cluster and similarly a few of the *Foxc1-KO* cells in the pia cluster were assigned by Seurat to the *Foxc1-KO* pia-like cluster. These were reassigned clusters based on their t-SNE space. A few cells were “undetermined” by Seurat FindClusters module. The neural, endothelial, monocyte, mural, osteogenic precursor and proliferating osteogenic precursor cells were assigned based on t-SNE space.

### Immunofluorescence and imaging

E14.5 embryos were collected and whole heads fixed overnight with 4% paraformaldehyde followed by 20% sucrose and frozen in OCT compound (Tissue-Tek). Tissue was cryosectioned in 12µm increments and tissue-mounted slides were subjected to antigen retrieval (except Cx43 antibody) by immersing the slides in 0.01M citric acid and heating in a pressure cooker for six minutes. The tissue was permeabilized by incubating for 10 minutes at room temperature in PBS with 0.1% Triton-X (Sigma) and blocked in 2% lamb serum/0.05% Triton-X solution for 40 minutes at room temperature. Incubation in the following primary antibodies was conducted overnight at 4°C in the blocking solution: rabbit anti-S100A6 (1:100; Novus NBP2-44492), rabbit anti-CRABP2 (1:100; Proteintech 10225), rabbit anti-Crym (1:100; Proteintech 12495), rat anti-Cx43 (1:1,000; Sigma C6219), rabbit anti-Col4 (1:200; BioRad 2150-1470) and chicken anti-GFP (Invitrogen A10262). Following incubation with primary antibodies, tissue sections were incubated for 60 minutes with appropriate Alexafluor-conjugated secondary antibodies (Invitrogen), Alexafluor 647-conjugated isolectin-B4 (1:100; Invitrogen I324550) and DAPI (1:1000; Invitrogen). Meninges E15.5 whole mounts were prepared and immunolabled using a protocol described in (Louveau, et al., 2018). In brief, brains were fixed overnight at 4°C in 2% paraformaldehyde and telencephalon meninges were removed and washed in PBS. Antibodies used in whole mount meninges staining were: rabbit anti-S100a6, rabbit anti-Raldh2 (1:200, Sigma HPA010022), rabbit anti-Col4 and rat anti-PECAM (1:100; BD Bioscience 557355). Confocal images were obtained using a Zeiss 780 Laser Scanning Microscope with associated Zeiss Zen software.

## Human fetal tissue collection and immunostaining

Human fetal tissue collection was obtained from a spontaneous abortion and after parents’ informed consent given to the Foetopathologie Unit at Hôpital Robert Debré and in accordance with French legislation. The gestational age (19wk) was based on first-trimester sonography crown-rump length measurement and confirmed at autopsy by the evaluation of fetal biometry and organ and skeletal maturations. Brains were removed and fixed in 4% buffered formalin added with 3 g/L of ZnSO_4_ for approximately 2 weeks before to be processed for paraffin embedding and sectioned at 6µm increments. Following de-parafinization, tissue staining, and imaging was performed as described for mouse tissue. In addition to above listed antibodies, we used mouse anti-µ-Crystallin (1:250; Invitrogen; cat# PIMA525192) in conjunction with antibodies to CRABP2 or S100a6.

## Acknowledgments

This work was supported by the National Institutes of Health/National Institute of Neurological Disorders and Stroke (R03 NS104566 and R01 NS098273 to J.A.S.). We thank members of the Siegenthaler and Jones’ labs for comments on the manuscript. We thank Damian Pawlikowski for his suggestion of the name ‘ceiling cell’ to describe a population of meningeal fibroblasts discovered as part of the single cell analysis of the meninges.

