## Supplemental Figures 1-7 for "A cellular atlas of the developing meninges reveals meningeal fibroblast diversity and function"

### Supp. Fig. 1

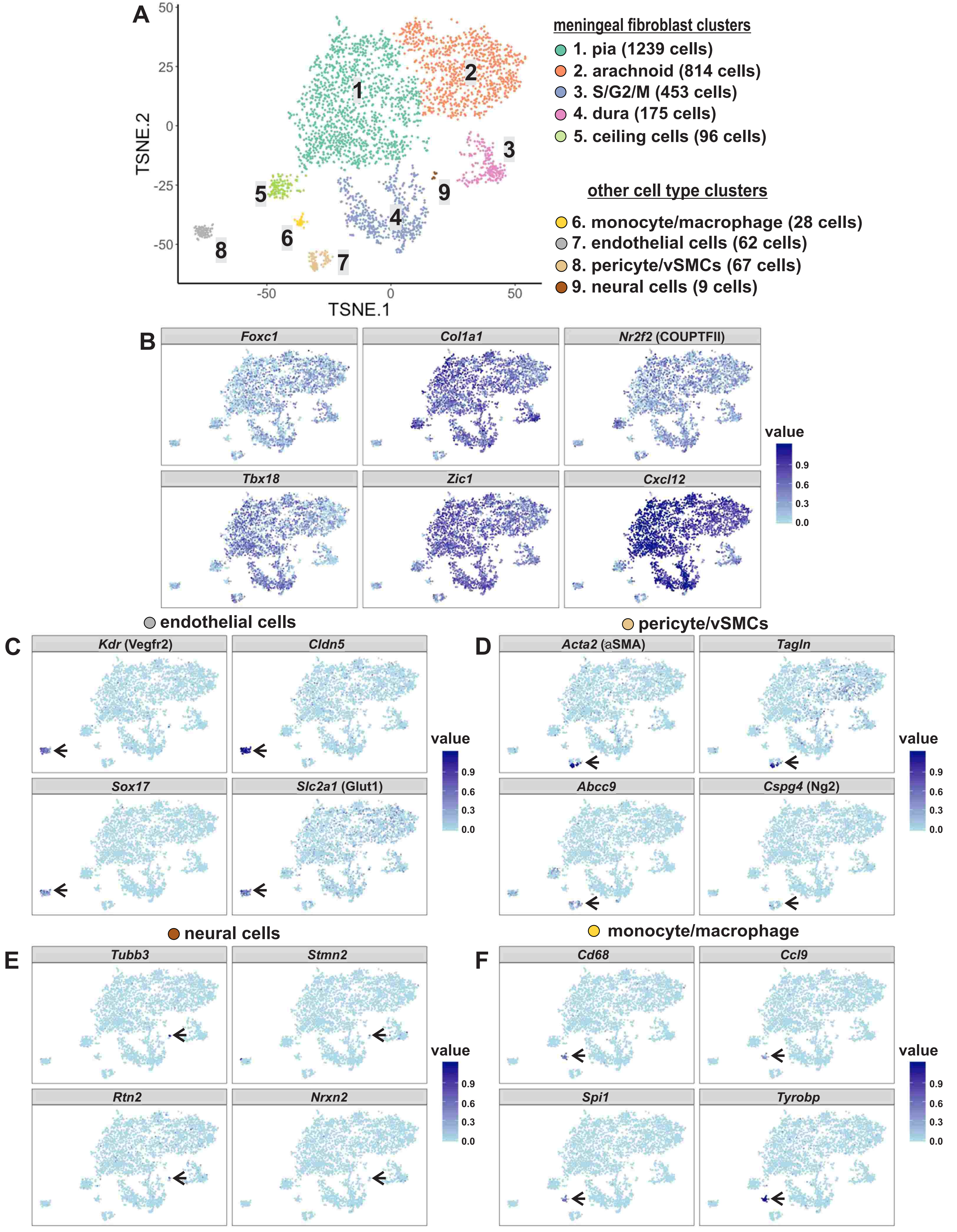
A
TSNE.2
TSNE.1
50
25
0
-25
-50
-50
0
50
1
2
3
4
5
6
7
8
9
meningeal fibroblast clusters
1. pia (1239 cells)
2. arachnoid (814 cells)
3. S/G2/M (453 cells)
4. dura (175 cells)
5. ceiling cells (96 cells)
other cell type clusters
6. monocyte/macrophage (28 cells)
7. endothelial cells (62 cells)
8. pericyte/vSMCs (67 cells)
9. neural cells (9 cells)
B
Foxc1
Col1a1
Nr2f2 (COUPTFII)
Tbx18
Zic1
Cxcl12
value
0.9
0.6
0.3
0.0
endothelial cells
C
Kdr (Vegfr2)
Cldn5
Sox17
Slc2a1 (Glut1)
pericyte/vSMCs
D
Acta2 (aSMA)
Tagln
Abcc9
Cspg4 (Ng2)
neural cells
E
Tubb3
Stmn2
Rtn2
Nrxn2
monocyte/macrophage
F
Cd68
Ccl9
Spi1
Tyrobp

**SUPPLEMENTAL FIGURE** 1 | Clustering and gene expression in all cells isolated from E14.5 *Col1a1-GFP+;Foxc1*$^{+/+}$ meninges. (A) Principal component analysis followed by 2D tSNE plotting of 2,943 cells from the control population (E14.5 embryo *Col1a1-GFP+;Foxc1*$^{+/+}$) comprising meningeal fibroblasts, monocytes/macrophages, endothelial cells, pericytes/vascular smooth muscle cells (vSMCs) and neural cells. (B) Expression of individual genes typically expressed in meningeal fibroblasts in tSNE space. (C) Expression in tSNE space of individual genes typically expressed in endothelial cells in tSNE space; arrow points to endothelial cell cluster. (D) Expression in tSNE space of individual genes typically expressed in pericytes/vSMCs in tSNE space; arrow points to pericyte/vSMC cell cluster. (E) Expression in tSNE space of individual genes typically expressed in neural cells in tSNE space; arrow points to neural cell cluster. (F) Expression in tSNE space of individual genes typically expressed in monocytes/macrophages in tSNE space; arrow points to monocyte/macrophage cell cluster.

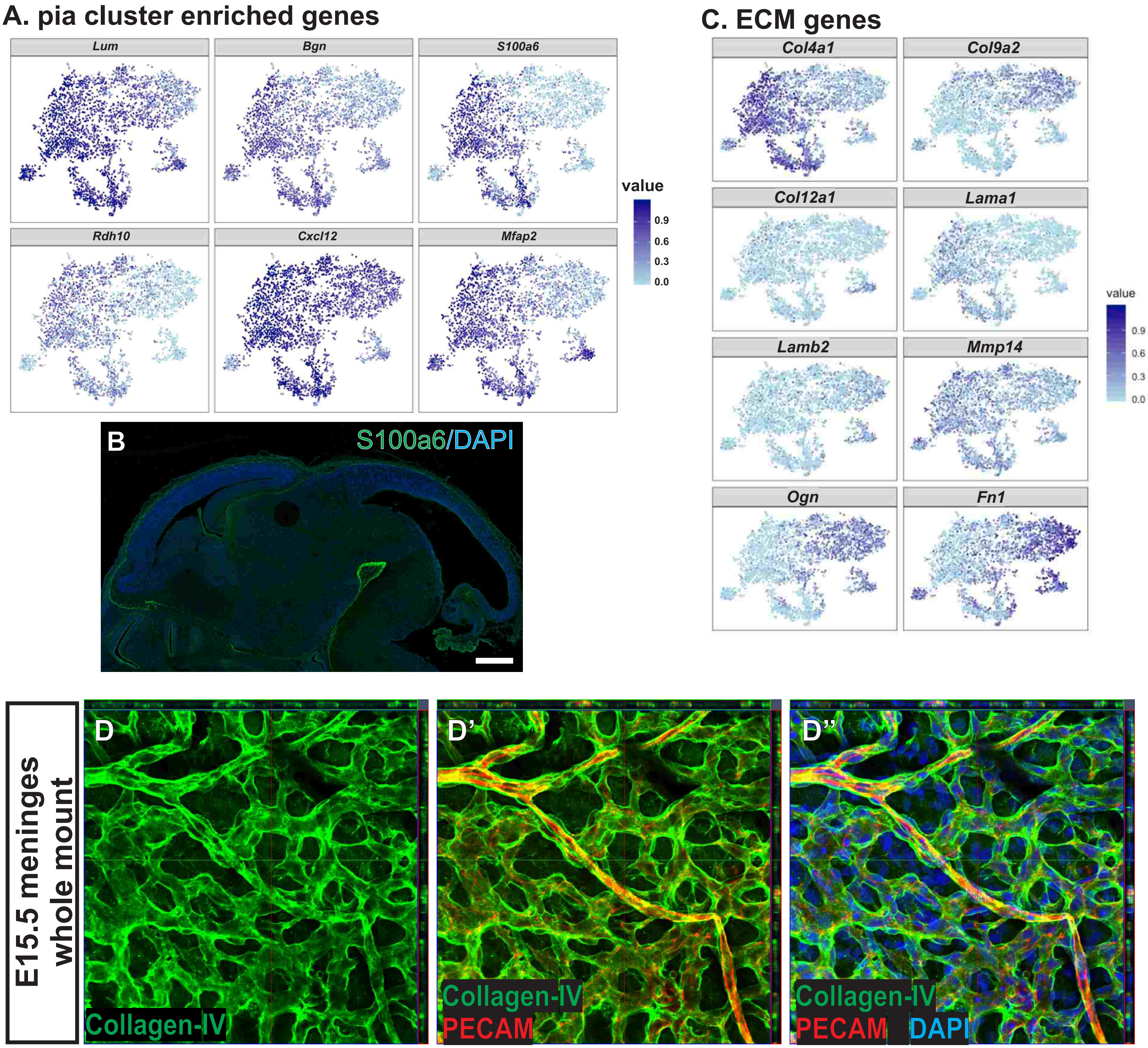


**SUPPLEMENTAL FIGURE 2** | (A) Expression of individual genes with enriched expression in the pia cluster shown in tSNE space. (B) Confocal image of immunofluorescence staining for S100a6 in sagittal section of E14.5 mouse embryo with DAPI to identify cell nuclei. (C) tSNE expression plots of ECM genes. (D) E15.5 meninges whole mount depicting Collagen-IV protein expression in the perineural vascular plexus within the meninges (labeled with PECAM) and in cells of the pial layer. Scale bar = 500 μm (B).

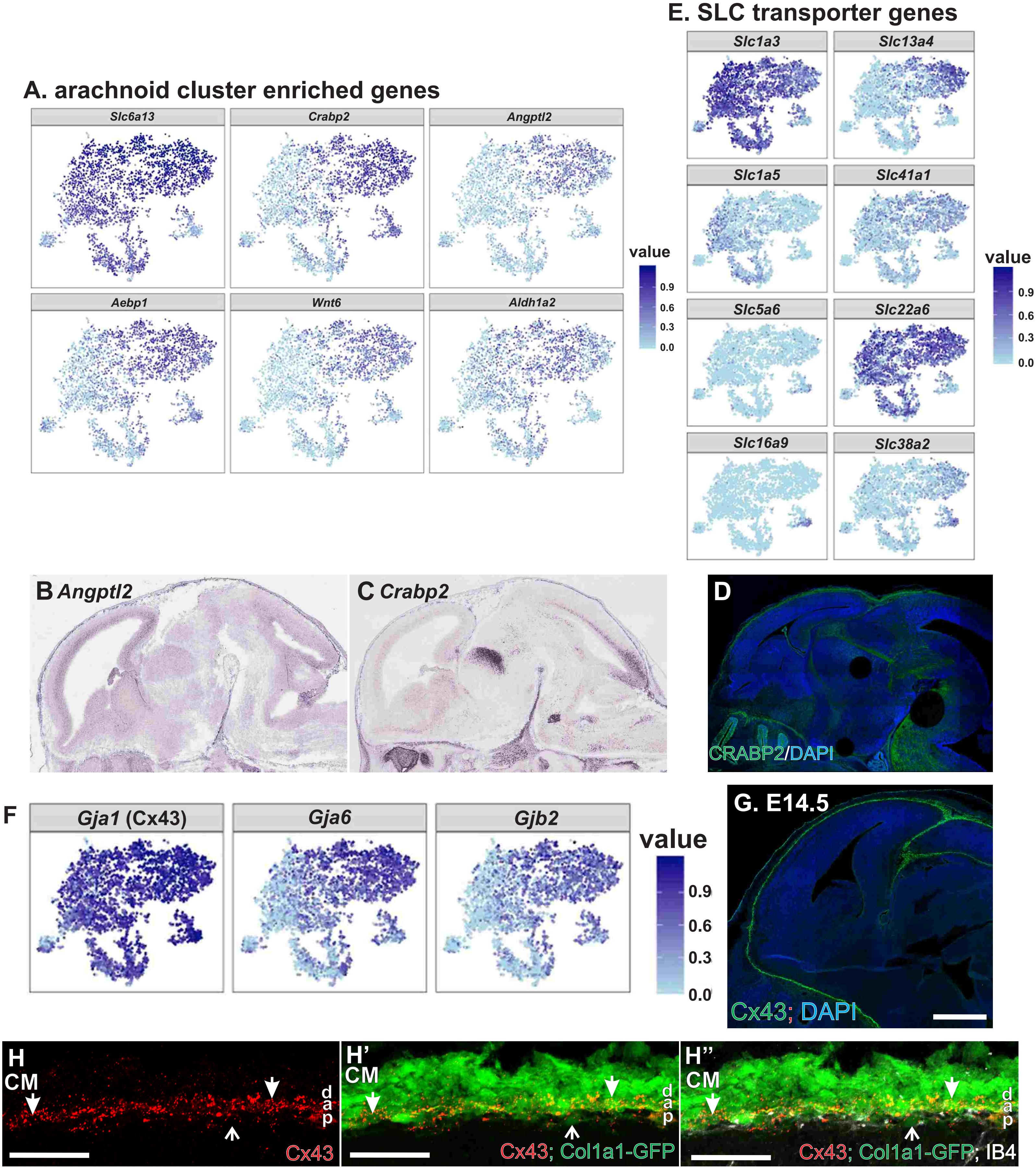


**SUPPLEMENTAL FIGURE 3** | (A) Expression of individual genes with enriched expression in the arachnoid cluster shown in tSNE space. (B, C) RNA in-situ hybridization showing *Angptl2* and *Crabp2* expression in E14.5 mouse embryo (Eurexpress.org). (D) Immunofluorescence of CRABP2 in sagittal section of E14.5 mouse embryo with DAPI to identify cell nuclei. (E) tSNE expression plots of SLC transporter genes. (F) tSNE expression plots of connexin genes. (G) Immunofluorescence of Cx43 in sagittal section of E14.5 mouse embryo with DAPI to identify cell nuclei. (H) Cx43 immunofluorescence in the meninges overlaying the neocortex in a E14.5 *Col1a1-GFP/+* embryo. Arrows indicated Cx43+/GFP+ cells in the arachnoid (a) and dura (d) and open arrows indicate Cx43+/GFP+ cells in the pia. Scale bars = 500 m (G) and 100 m (H).

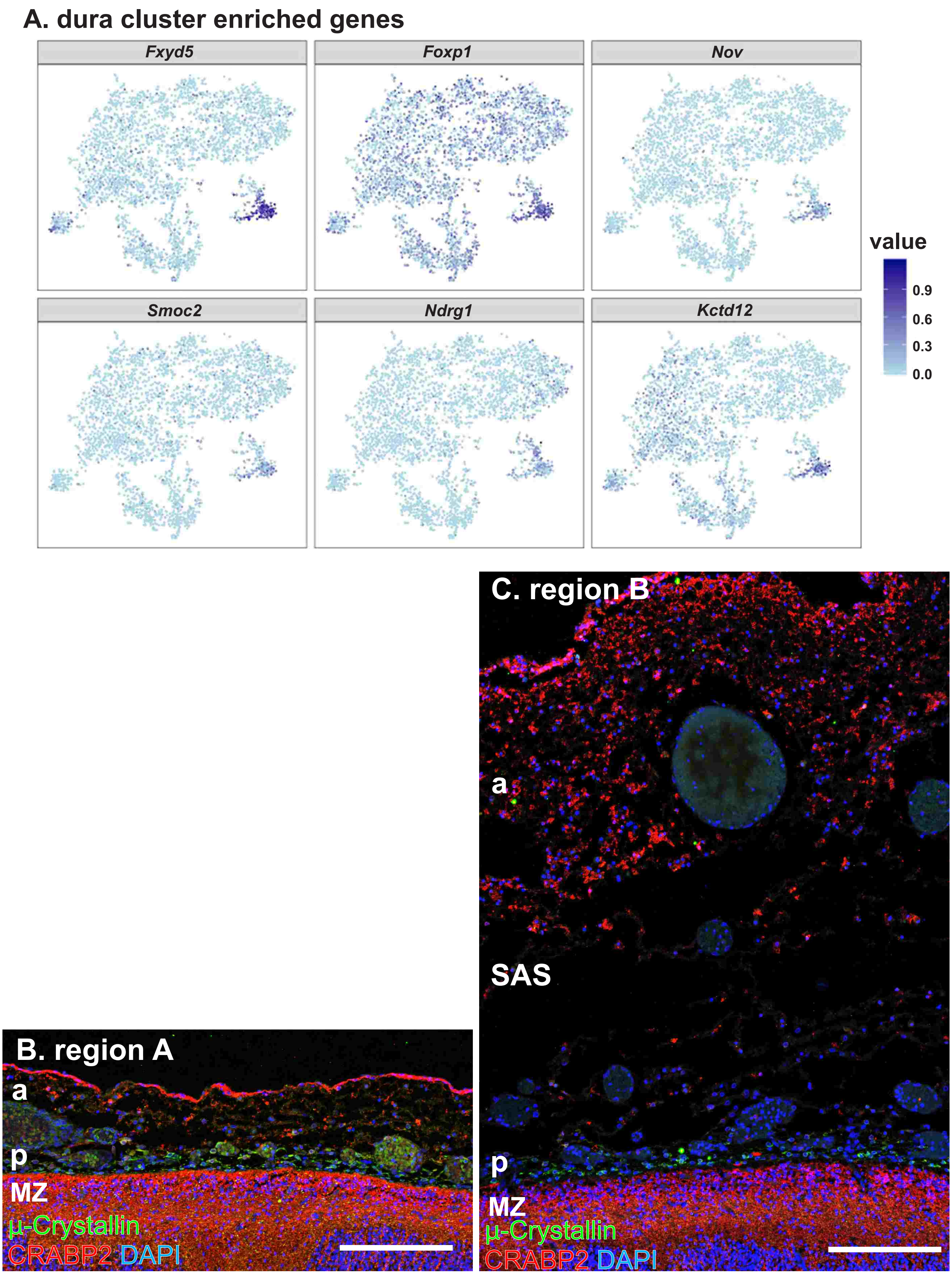


**SUPPLEMENTAL FIGURE 4** | (A) Expression plots of individual genes with enriched expression in the dura cluster shown in tSNE space. (B) CRAPB2 and -Crystallin+ immunolabeling in region A and region B of the human fetal meninges. SAS= subarachnoid space, p=pia, a=arachnoid, mz=marginal zone. Scale bars = 200 m (B, C).

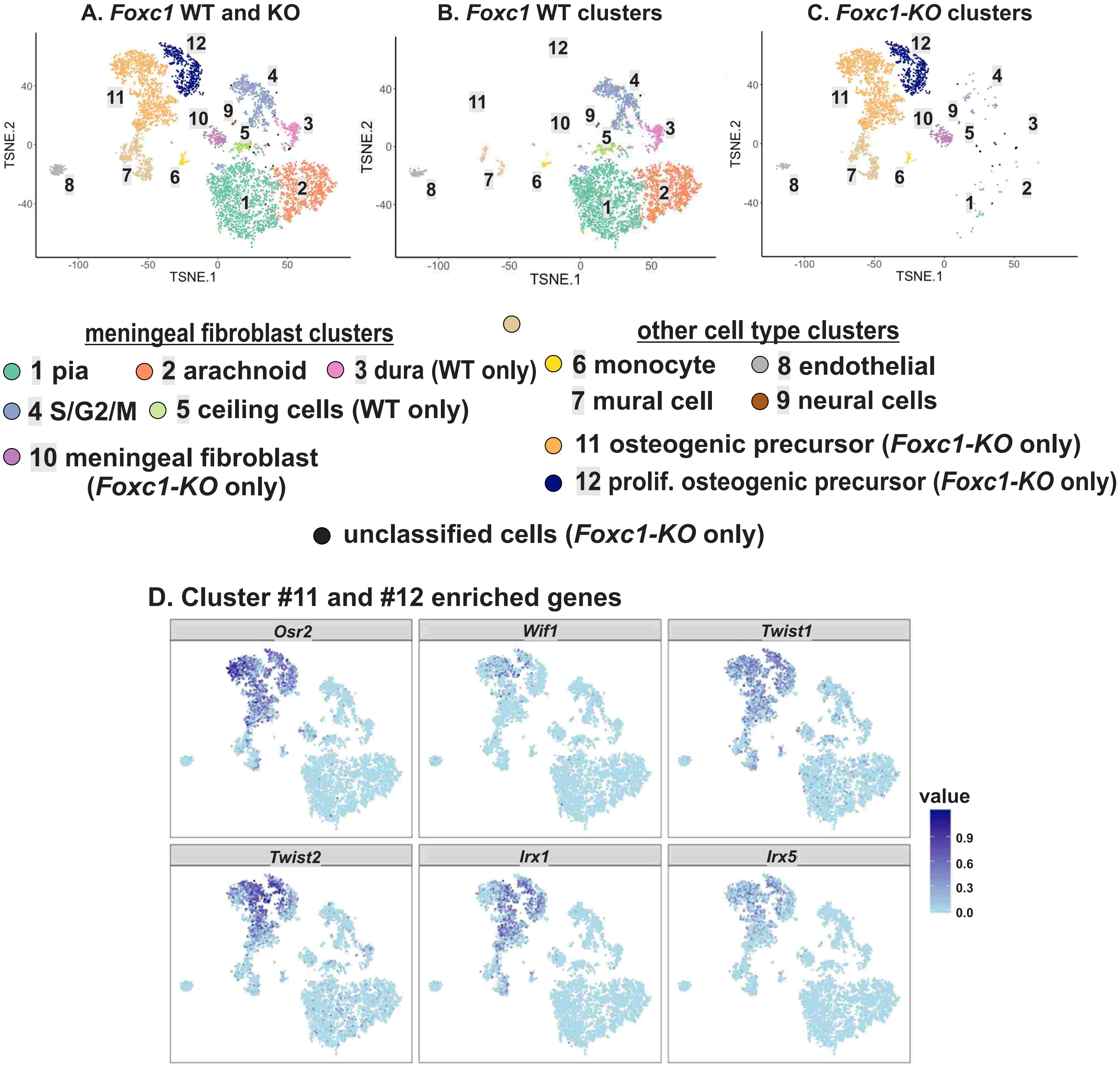


**SUPPLEMENTAL FIGURE** 5 | Cluster designation for combined analysis of E14.5 control (*Col1a1-GFP+;Foxc1*$^{+/+}$) and *Foxc1-KO* (*Col1a1-GFP;Foxc1*$^{lacZ/lacZ}$) cells. (A-C) tSNE plot of control and *Foxc1-KO* cell clusters together in a single plot (A) and in individual plots (B-C). Clusters 10, 11, 12 contain only *Foxc1-KO* cells. (D) Expression in tSNE space of individual genes enriched in clusters #11 and 12 that contain only *Foxc1-KO* cells.

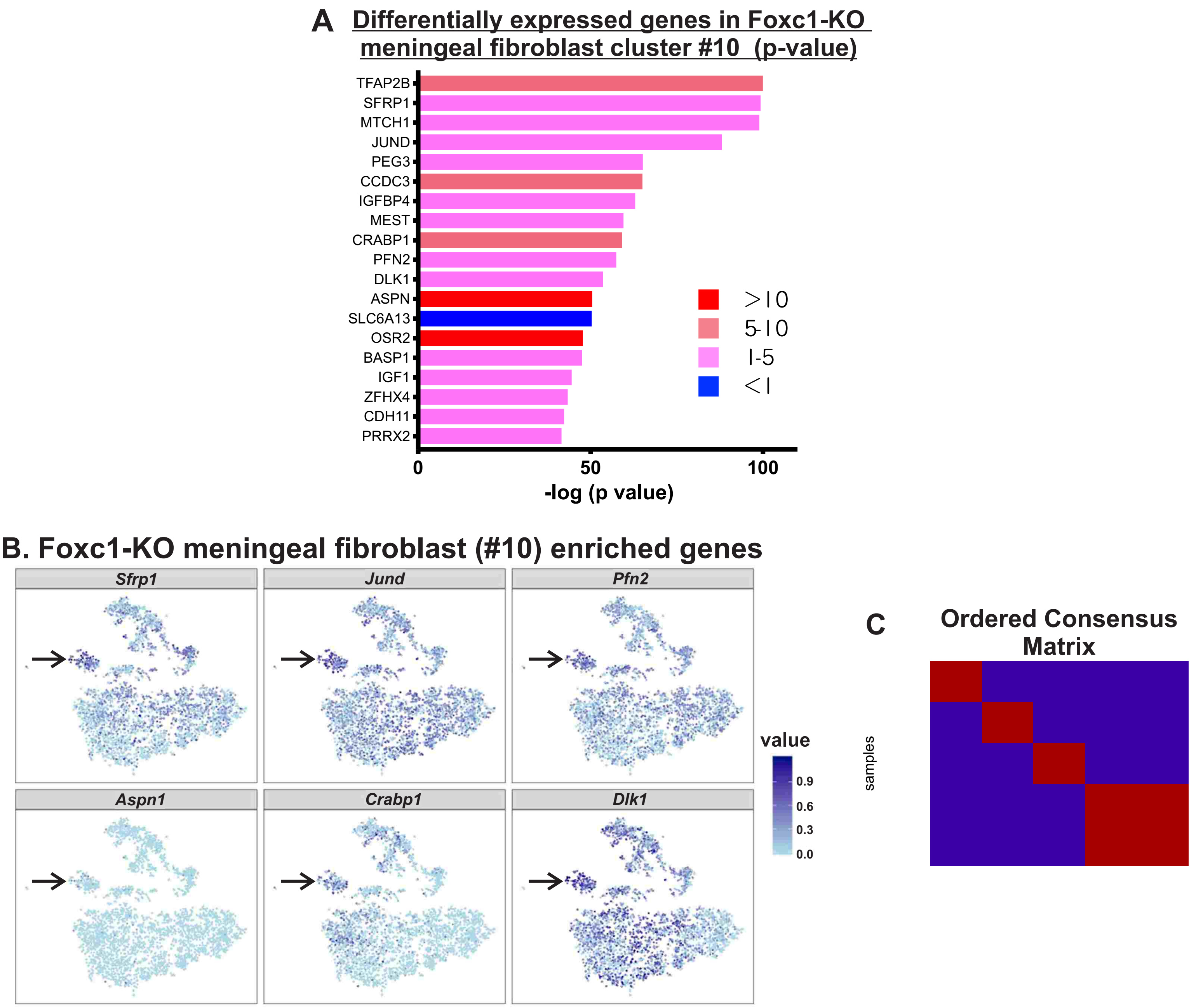


**SUPPLEMENTAL FIGURE** 6 | Expression of genes enriched in *Foxc1-KO* fibroblasts cluster #10. (A) Bar graph depicts genes with greatest enrichment (ER 1-5) or depletion (ER <1) in the *Foxc1-KO* meningeal fibroblast cluster (#10). Bar length represents the negative base-10 log of p value and color represents ratio of each gene's mean expression in cluster #10 versus all other control meningeal fibroblasts (ER). (B) Expression of individual genes with enriched expression in the *Foxc1-KO* meningeal fibroblast (#10) in tSNE space; arrow points to *Foxc1-KO* cluster #10. (C) Ordered consensus matrix for four subgroups (k=4) from NMF analysis depicted in Fig. 6E; cophenetic correlation in the NMF analysis was maximized for 4 subgroups (k=4, coph = 1).

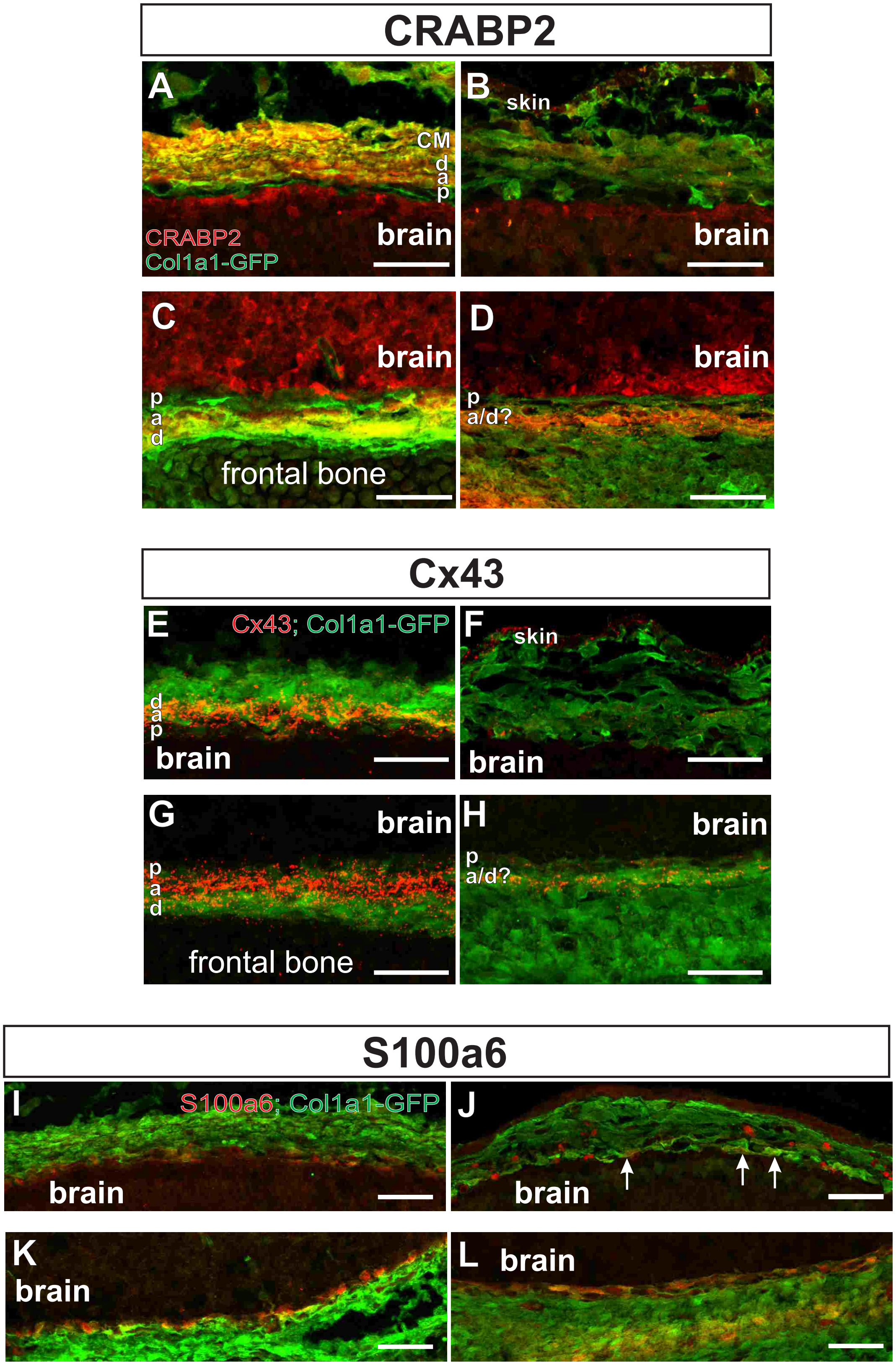


**SUPPLEMENTAL FIGURE 7** | (A-D) CRABP2 depicted in Fig. 7C-F with *Col1a1-GFP* label to identify fibroblasts. (E-H) Cx43 depicted in Fig. 7I-L with *Col1a1-GFP* label to identify fibroblasts. (I-L) S100a6 depicted in Fig. 7O-R with *Col1a1-GFP* label to identify fibroblasts (arrows indicate S100a6+/GFP+ cells in the pia). Scale bars = 100 m. a=arachnoid, d=dura, p=pia, cm=calvarial mesenchyme.
